# A spatially continuous cell-type atlas reveals multiscale organization of the human cortex

**DOI:** 10.64898/2026.11.10.765231

**Authors:** Wenkun Lei, Xiaohan Tian, Changsheng Dong, Jing Lou, Xinghui Zhao, Xinyi Dong, Lin Du, Ruoxin Yang, Yuxuan Hong, Yu Yu, Chen Yang, Linna Qian, Shu Liu, Ang Li, Meng Wang, Bing Liu

## Abstract

Neuroimaging has increasingly revealed fine-grained spatial heterogeneity within the human cortex, uncovering regional differentiation beyond conventional cortical divisions. However, understanding the cellular architecture underlying these imaging features requires spatially continuous cell-type references that capture both cortex-wide organization and regional specialization. Here, we introduce RIDGE (Regional Integration of Deconvolution with Guided Estimation), a deconvolution framework that anchors multiple regional references to continuous cortex-wide transcriptomic maps, thereby recovering cortex-wide organizational patterns while supporting region-specific cell-type inference. RIDGE reconstructs spatially continuous distributions of 24 major cortical cell types across the human cortex. The resulting maps are robust across analytical pipelines and independent reference datasets, recover established cell-type associations with known cortical features, and recapitulate the canonical primary sensory–allocortical/periallocortical (Pr–Al) axis of cortical cellular organization. At finer scales, RIDGE resolves distinct cellular profiles across default mode, motor, and visual systems, revealing local cellular specialization within each system. By relating these cellular maps to cortical imaging features, we demonstrate their utility for investigating cellular contributions to global cortical organization and regional specialization. Together, RIDGE provides a unified framework and spatially continuous cellular reference resource for integrating cellular architecture with human cortical imaging across spatial scales.

## Introduction

The human cerebral cortex exhibits remarkable structural and functional organization, shaped by the diversity, distribution and spatial arrangement of neuronal and non-neuronal cell types. Recent advances in single-cell and single-nucleus transcriptomics have greatly expanded our understanding of cellular and molecular diversity in the human brain, revealing distinct cell types and transcriptional programs across cortical regions^1–3^. In parallel, multimodal neuroimaging provides spatially continuous views of cortical structure, neural dynamics and functional organization across the whole brain^4–6^. Bridging these complementary scales could extend cellular information to the continuous spatial scale of macroscale brain maps, enabling a more complete understanding of how cellular architecture supports cortex-wide organization.

Recent studies have sought to link cell-type-associated transcriptional signatures with macroscale cortical and neuroimaging phenotypes^7–9^. However, translating cellular information from discrete reference samples into continuous cortical maps remains challenging. Single-cell sequencing datasets provide rich cellular specificity but limited anatomical coverage, whereas cortex-wide transcriptomic maps offer broad spatial coverage without direct cellular resolution. Existing strategies typically rely on either single-region or pooled cellular references, yet neither can simultaneously support robust cortex-wide inference and preserve regional transcriptional specificity. A central question is therefore whether region-specific cellular information can be incorporated into cortex-wide inference while retaining comparability across cortical regions.

Here, we develop RIDGE (Regional Integration of Deconvolution with Guided Estimation), an integrated deconvolution framework that unifies regionally sampled single-nucleus RNA-sequencing references^1,2^ with spatially distributed transcriptomic data from the Allen Human Brain Atlas^10^, enabling cortex-wide cellular estimation while preserving regional specificity. RIDGE uses the cortex-wide transcriptomic maps as a common anchor to harmonize regional single-nucleus references and derives region-specific cell-type signatures for reconstructing spatially continuous distributions of 24 major cortical cell types^11^. We assess the stability across analytical pipelines and independent reference datasets and validate their biological plausibility against established cell-type-specific features of cortical organization. The resulting maps robustly capture the Pr–Al organization of cortical cell-type composition^12^. They further resolve distinct cellular profiles across the default mode network (DMN), motor and visual systems, as well as finer spatial differentiation within each system. Together, these analyses demonstrate the ability of the framework to recover biologically meaningful cellular organization across cortex-wide, system-level, and local spatial scales.

## Results

### RIDGE framework overview

RIDGE is a transcriptomic deconvolution framework that uses cortex-wide transcriptional anchoring and regional reference integration to reconstruct spatially continuous cell-type maps across the human cortex. The core methodological steps are outlined below, with full details provided in the **Methods** (**Fig. 1a**). **(1)** Transcriptomic inputs (**Fig. 1a**): We used AHBA-derived Dense Expression Maps (DEMs) as the transcriptomic target for deconvolution, preserving fine-grained anatomical detail and spatial continuity across the cortex^13^; we partitioned the multi-region single-nucleus datasets by Brodmann area to generate region-specific references that retain regional transcriptional variation (**Supplementary Fig. 1; Supplementary Table 1**). **(2)** Regional reference anchoring (**Fig. 1a**): To harmonize regional references while reducing region-dependent scaling effects, each regional single-nucleus reference was concatenated with the cortex-wide DEM target and jointly normalized. The cortex-wide transcriptomic target served as a common anchor for regional references, from which region-specific cell-type markers were identified for subsequent region-matched deconvolution. **(3)** Region-matched deconvolution (**Fig. 1a**): To preserve regional specificity during inference, each regional reference was deconvolved against the transcriptomic target from its corresponding Brodmann area, yielding vertex-wise cell-type estimates for that region. **(4)** Geometry-constrained cortical field reconstruction (**Fig. 1a**): To integrate region-wise estimates within the intrinsic geometry of the cortical surface, we mapped them to their corresponding cortical territories and applied a surface-constrained spatial operator to reconstruct continuous cortex-wide cell-type fields in fsLR-32k space.

**Fig. 1.**
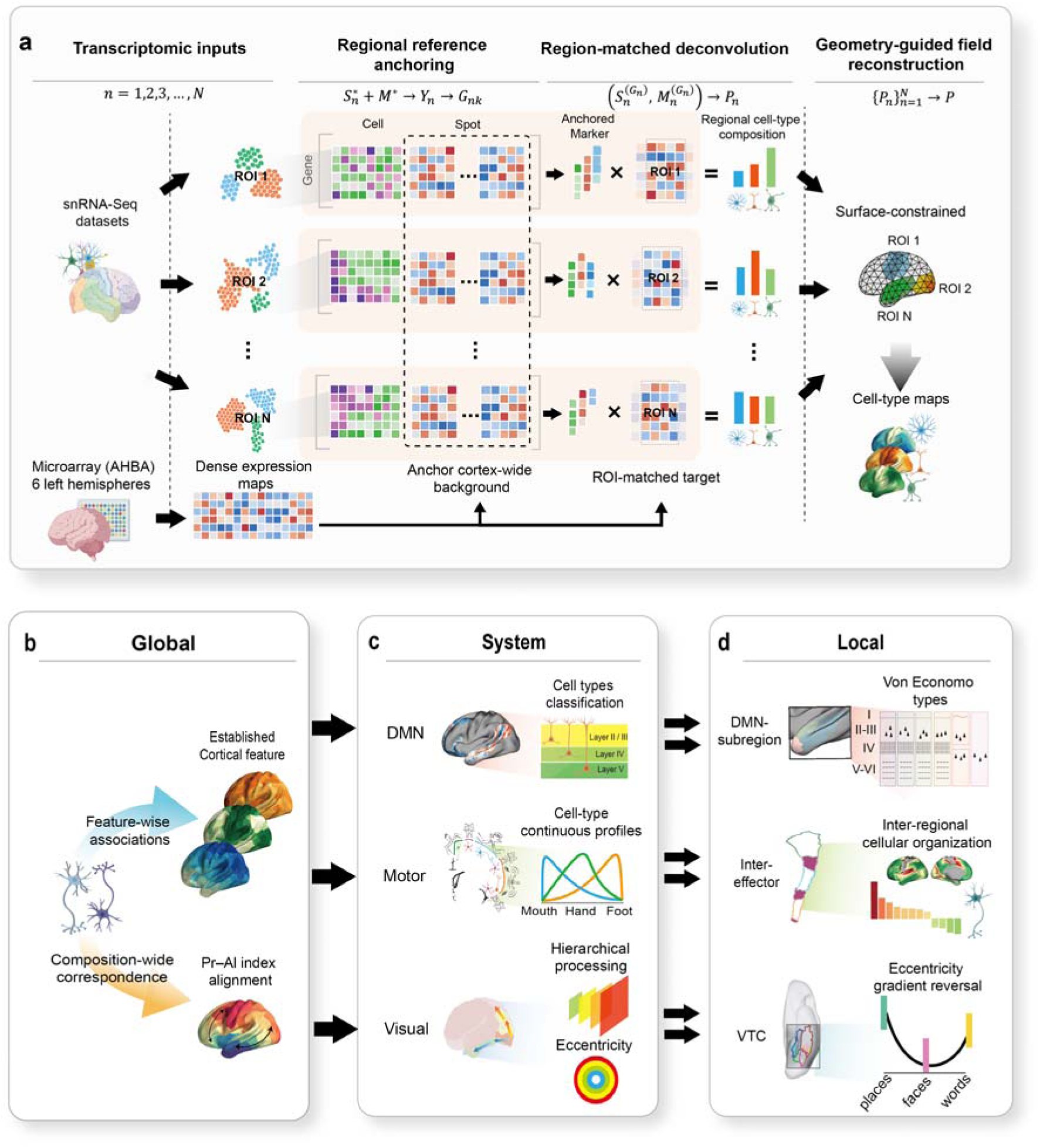
A multi-region deconvolution framework for reconstructing cortical cell-type organization across spatial scales. **(a)** Workflow of RIDGE. AHBA-derived DEMs provide the cortex-wide transcriptomic target M* (genes × vertices), while multi-region single-nucleus data are partitioned into Brodmann-defined references *S*\**_n_*(genes × cells). Each *S*\**_n_* is jointly normalized with M* to generate the anchored matrix Y_n_, from which cell-type-specific marker sets G_nk_ are identified. The resulting reference and markers are then applied to the ROI-matched target M_n_ to estimate regional cell-type composition P_n_. Estimates across all regions are integrated within the cortical surface geometry to reconstruct continuous cortex-wide cell-type maps. **(b)** Global-scale evaluation of the framework-derived cell-type maps. Feature-wise analyses relate individual cell-type maps to established cortical features. Composition-wide analyses assess the alignment between the dominant pattern of cortical cell-type composition and the Pr–Al organization. **(c)** System-scale analyses in three representative cortical systems. Within the DMN, cell-type maps are related to microstructural differentiation; within the primary motor cortex, cell-type variation is mapped along the proximal–distal axis; and within the visual cortex, cellular variation is examined in relation to hierarchical processing and eccentricity. **(d)** Local-scale analyses within each system. Cell-type organization is evaluated across histologically differentiated DMN subregions; within inter-effector regions of the primary motor cortex; and across cytoarchitectonic and category-selective organization in the ventral temporal cortex (VTC). Certain schematic elements in this figure were created with **biogdp.com**^14^.

We next evaluated the technical robustness and regional reference sensitivity of RIDGE before characterizing the reconstructed cellular organization at global (**Fig. 1b**), system (**Fig. 1c**) and local (**Fig. 1d**) scales.

### Generation and validation of RIDGE-derived cortex-wide cell-type maps

We implemented RIDGE using a standardized reference taxonomy comprising 24 cortical cell types and evaluated RIDGE-derived maps using complementary technical and biological benchmarks (**Fig. 2a; Supplementary Tables 2–3**). The spatial distributions of cell types were robust across alternative cortical transcriptomic processing pipelines and reproducible across independent single-nucleus reference datasets (**Supplementary Fig. 2a, b**). Cross-reference validation further showed that cell types with greater regional transcriptional variability exhibited lower spatial consistency when inferred from different cortical references (rho = −0.50, *P* = 0.01; **Supplementary Fig. 3a–c** and **Supplementary Table 4**). Together, these analyses indicate that RIDGE generates stable and biologically interpretable estimates of relative cellular variation while reflecting regional variation in reference transcriptional profiles. We next examined whether the maps preserved fine-scale cellular variation within well-characterized cortical systems. In primary motor cortex (area 4, defined by the HCP-MMP1 atlas), the peak of Oligo values along the ventral–dorsal position coincided with the peak of myelin values (**Fig. 2b**). In the visual system, L4 IT neurons showed strong enrichment in V1 that gradually declined toward higher-order visual regions (V2–V4; **Fig. 2c**), mirroring the transition from sensory input to integrative processing^15,16^. These findings indicate that the framework captures fine-grained cellular differentiation consistent with known cortical organization.

**Fig. 2.**
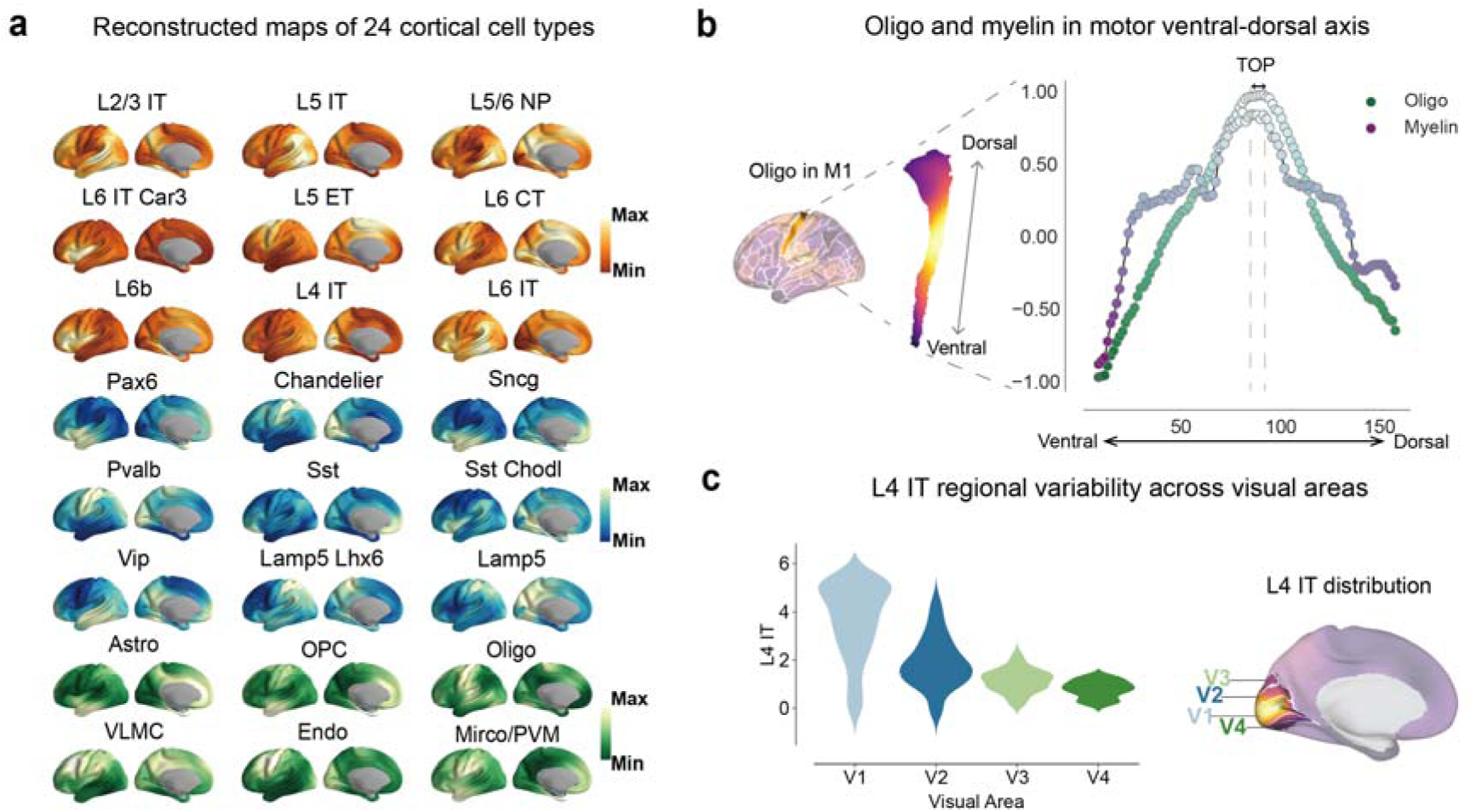
RIDGE-derived cell-type maps and regional cellular organization. **(a)** Cortical maps of 24 RIDGE-derived cell types. Surface maps show vertex-wise distributions reconstructed by RIDGE. Colors indicate major cell classes: excitatory neurons (orange), inhibitory neurons (blue), and non-neuronal cells (green). **(b)** Oligo and myelin along the ventral–dorsal position of the primary motor cortex (defined by HCP-MMP1 area 4). Values were normalized to [−1, 1], and dashed lines indicate peak positions. **(c)** Regional differentiation of L4 IT across early visual areas (V1–V4) defined by the HCP-MMP1 atlas.

### RIDGE-derived cell-type maps reveal global principles of cortical cellular organization

We first evaluated four prespecified associations grounded in well-established biological links between specific cell types and cytoarchitectonic, evolutionary, functional, and neurophysiological features of the cortex^1,17–21^. A composite L2/3 IT–L5 IT measure was positively associated with evolutionary cortical expansion (*r* = 0.43, *P*_spin_ = 0.003; **Fig. 3a**). L4 IT distributions showed strong spatial correspondence with layer 4 thickness (*r* = 0.66, *P*_spin_ < 0.001; **Fig. 3a**), and the OPC-to-oligodendrocyte ratio correlated positively with the principal functional connectivity gradient (*r* = 0.54, *P*_spin_ < 0.001; **Fig. 3a**). Pvalb neuron distributions were negatively associated with high-gamma power (*r* =-0.68, *P*_spin_ = 0.018; **Fig. 3a**). These convergent associations across independent domains support the biological plausibility of the reconstructed cellular maps.

**Fig. 3.**
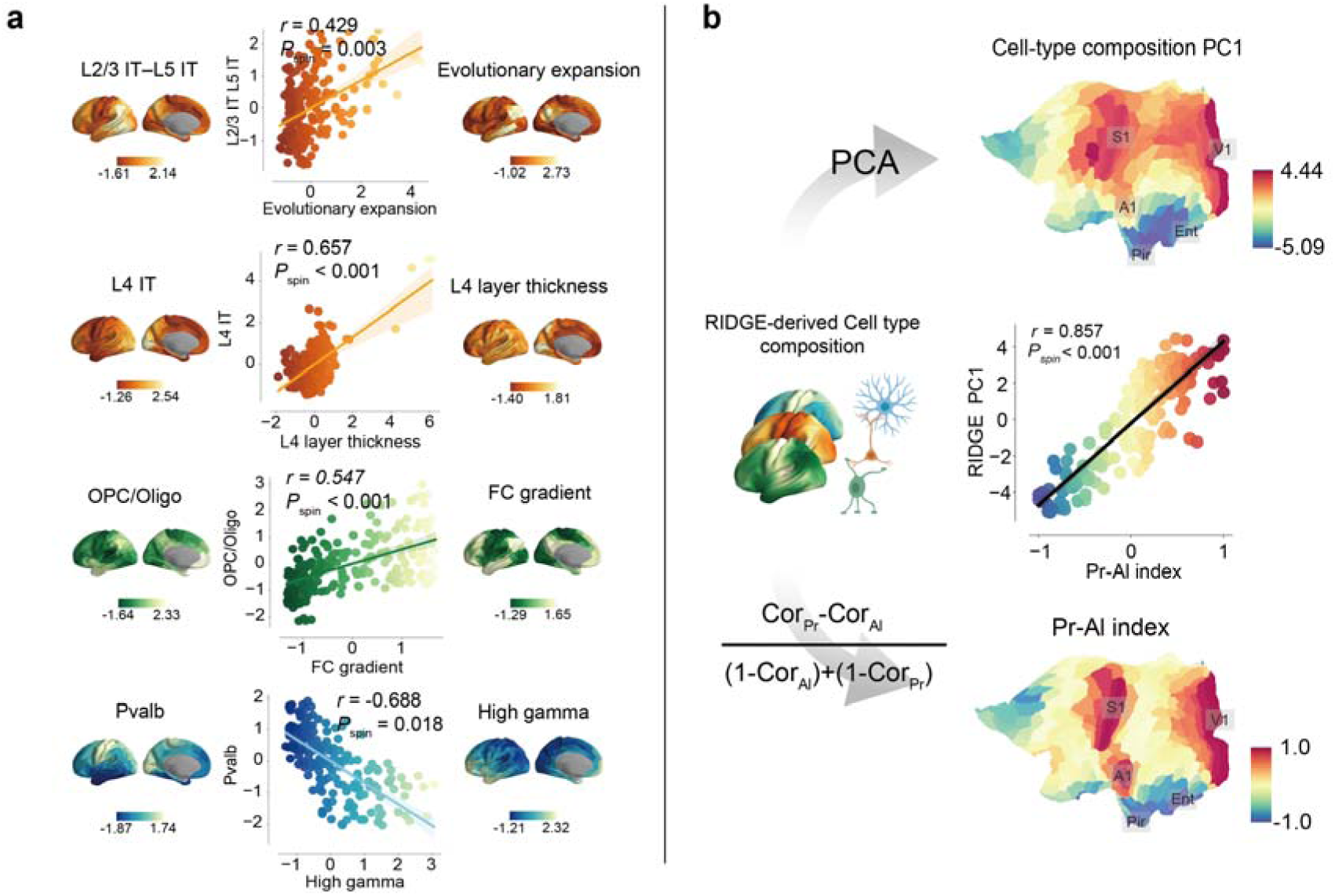
Global organization of RIDGE-derived cell-type maps. **(a)** Associations between selected cellular measures and established cortical features, including evolutionary expansion, layer 4 thickness, the principal functional connectivity gradient, and high-gamma power. Statistical significance was assessed using spin permutation testing (n = 1,000; shading indicates 95% confidence intervals). **(b)** PCA of the 24 RIDGE-derived cell-type maps identified a dominant spatial pattern of cortical cell-type composition that closely aligned with the Pr–Al organization (r = 0.857, *P*_spin_ < 0.001)

We then examined whether cortex-wide variation in cellular composition captured a major organizational pattern of cortical cell-type composition. Principal component analysis (PCA) of the 24 cell-type maps identified a dominant spatial mode that closely aligned with the Pr–Al organization of cortical cell-type composition (*r* = 0.857, *P*_spin_ < 0.001; **Fig. 3b**, **Supplementary Fig. 4a**)^12^. This organization was further supported by directly measured regional cell-type compositions in humans (**Supplementary Fig. 4b**). Comparable Pr–Al organization was also observed in macaque and marmoset datasets (**Supplementary Fig. 4c**). These findings indicate that the dominant cellular pattern is consistent with conserved principles of cortical organization. We next compared the spatial recovery of the Pr–Al pattern across reference strategies using geodesic deviation and spatial overlap. Across increasing Pr–Al tail thresholds, RIDGE consistently showed lower geodesic deviation and greater spatial overlap than the Lake visual, Lake frontal, and cross-area robust-marker strategies (**Supplementary Fig. 5a–c, Supplementary Tables 5–6**), indicating improved recovery of the spatial organization of the Pr–Al pattern. We further used the maps in cross-modal analyses of cortical molecular and disease-related organization, identifying systematic associations with cortical abnormalities across 13 neuropsychiatric disorders and regional neurotransmitter receptor distributions (**Supplementary Fig. 6, Supplementary Table 7**)^22,23^.

Together, these findings indicate that RIDGE-derived cellular maps capture reproducible global principles of cortical organization and provide a biologically informative representation for linking cellular composition with molecular, functional and disease-related cortical phenotypes.

### Cell-type–microstructure correspondence reveals DMN organization across scales

We next examined whether they captured within-system microstructural differentiation in the DMN, a transmodal system characterized by pronounced cytoarchitectural heterogeneity^24,25^. We systematically assessed the correspondence between the BigBrain primary gradient (G1) and all 24 cell-type maps using spin permutation tests with false discovery rate (FDR) correction (**Supplementary Fig. 7a**) ^26,27^. Three intratelencephalic neuron types—L2/3 IT, L4 IT, and L5 IT—showed significant positive correlations with G1 and were retained for subsequent analyses ( L2/3 IT: *r* = 0.36, *P*_spin_ = 0.048; L4 IT: *r* = 0.46, *P*_spin_ = 0.012; L5 IT: *r* = 0.44, *P*_spin_ = 0.031, **Fig. 4a**). These IT populations are major components of corticocortical projection pathways, consistent with their spatial association with DMN microstructural organization^28,29^. This correspondence was further supported by higher mean navigation efficiency in DMN regions enriched in these IT neuron types (**Supplementary Fig. 7b**)^30^.

**Fig. 4.**
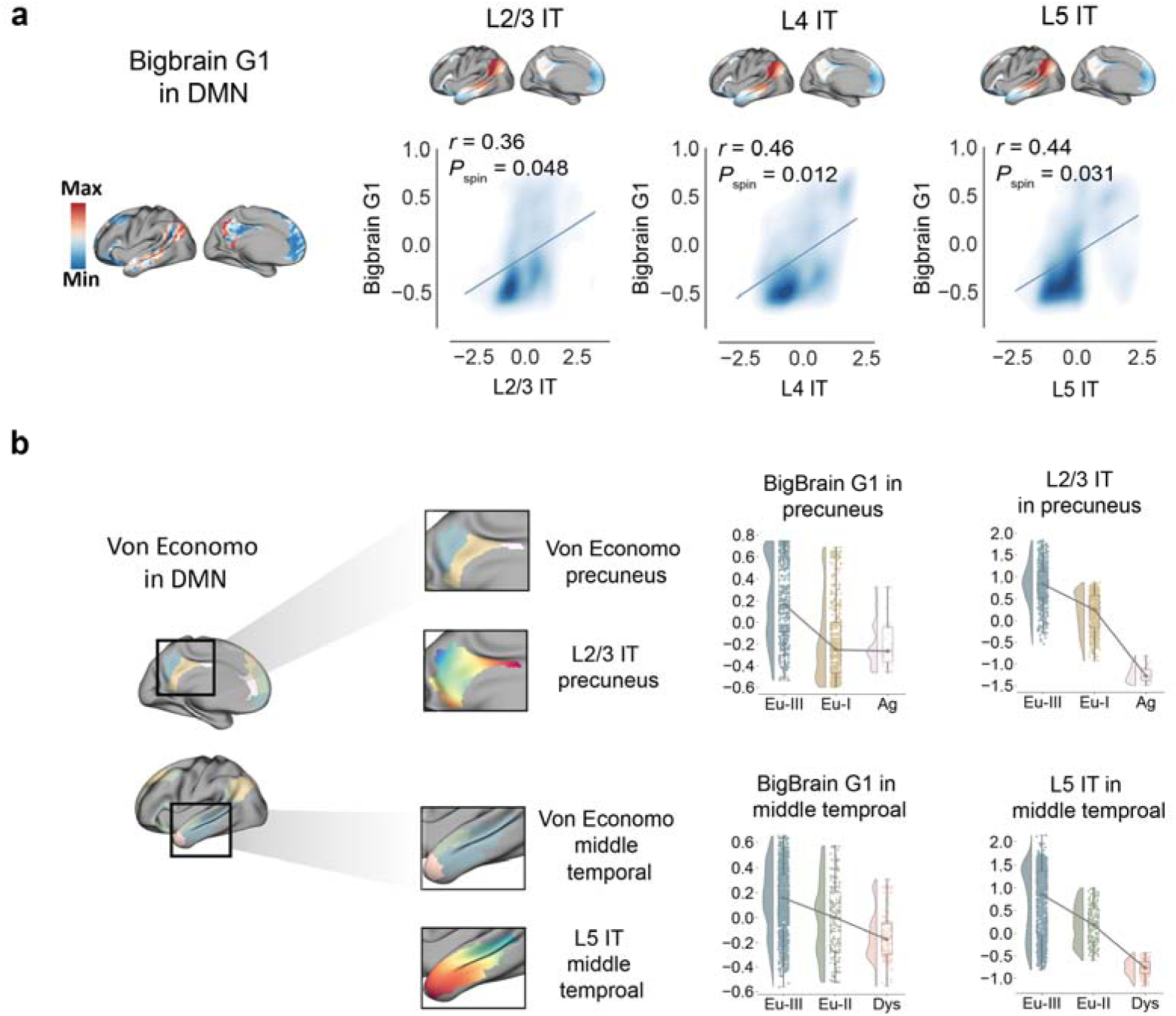
RIDGE-derived cell-type maps capture system-and local-scale microstructural differentiation within the DMN. **(a)** Density plots show the correspondence between BigBrain G1 and representative L2/3 IT, L4 IT, and L5 IT within the DMN (spin permutation, n = 1,000; shading: 95% CI). **(b)** Raincloud plots show the alignment between representative cell-type maps and microstructural gradients across DMN subregions. DMN subregions were parcellated according to Von Economo cytoarchitectonic classes. See also **Supplementary** Fig. 7c.

Moving from the system to the local scale, we further evaluated whether the inferred cell-type maps captured finer-grained histological differentiation across DMN subregions using the Von Economo cortical classification^31^. In the precuneus, G1 varied systematically across Von Economo cortical types, decreasing from Eu-I/II to Eu-I and Ag, with L2/3 IT showing a matching trend. A similar correspondence was observed in the middle temporal gyrus, where both G1 and L5 IT reached their highest values in Eu-III regions and progressively declined toward dysgranular cortex (**Fig. 4b** and **Supplementary Fig. 7c**). These concordant patterns indicate that the reconstructed cell-type maps capture histological variation at a finer anatomical scale within DMN subregions. In the prefrontal subregion, the correspondence between G1 and the reconstructed cell-type maps was preserved; however, neither followed the conventional transition from highly laminated to less differentiated cortex (**Supplementary Fig. 7c**), consistent with the distinctive cytoarchitectural organization of the prefrontal DMN^24,25^. This regional deviation further indicates that the framework preserves anatomically meaningful variation rather than imposing a uniform microstructural hierarchy across the cortex.

### Cellular organization captures motor topography and inter-effector functional organization

We next evaluated whether the reconstructed cell-type maps captured cellular specialization within the primary motor cortex (M1), a representative sensorimotor system with well-characterized somatotopic organization^32^. At the system level, we first examined cell-type distributions along the proximal–distal axis of M1, operationalized along the ventral–dorsal extent of area 4. We used HCP-MMP1 area 4 as the anatomical scaffold and Gordon’s parcellation to define effector-specific motor regions (mouth, hand, and foot) ^32^. Along this axis, distinct cell types showed ordered spatial gradients that recapitulated known features of M1 organization (**Fig. 5a**; **Supplementary Fig. 8a**). L5 ET neurons showed a clear decreasing trend from ventral to dorsal M1, whereas Oligo maps peaked in the hand region and Sst interneurons showed an increasing trend along the same axis (**Fig. 5a**). These spatial patterns were consistent with established features of M1 organization^33,34^, supporting the biological plausibility of the reconstructed cell-type maps.

**Fig. 5.**
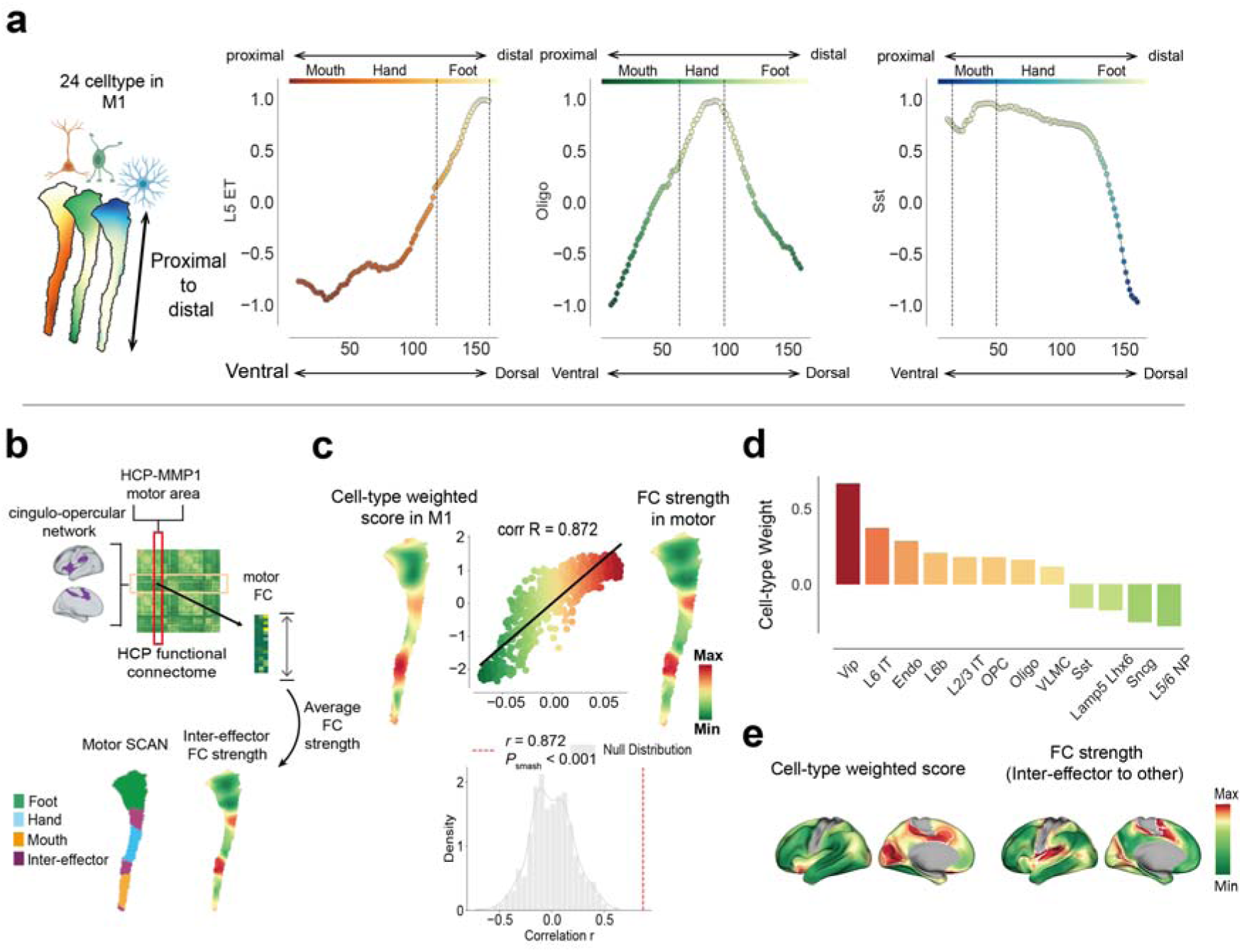
Cell-type maps reveal the functional topography of the motor cortex. **(a)** Distributions of L5 ET, Oligo, and Sst from RIDGE-derived cell-type maps along the proximal–distal axis of M1, defined using HCP-MMP1 area 4. Cell-type values were normalized to [−1, 1], and vertical dashed lines mark the mouth, hand, and foot representations defined by Gordon’s parcellation, corresponding to local peaks in the cellular profiles. **(b)** Workflow for deriving the inter-effector FC map between M1 and CON. **(c)** Spatial correspondence between the CCA-derived cell-type weighted map and inter-effector FC strength. Top, surface maps and vertex-wise scatter plot showing a strong spatial correlation (*r* = 0.872; *P*_smash_ < 0.001). Bottom, null distribution of correlation coefficients from spatial permutation testing. **(d)** CCA-derived weights of 12 nonredundant cell types. **(e)** The CCA-derived cellular signature and inter-effector-to-other FC strength exhibit overlapping spatial patterns, highlighting convergence in the aPFC and visual border regions (*P*_smash_ < 0.001). White outlines delineate the top 10% of values for each map.

At the local scale, we next focused on inter-effector regions embedded within the proximal–distal organization of M1. Based on previous work showing that these regions are preferentially coupled to the somato-cognitive action network (SCAN)^32^, we quantified inter-effector functional connectivity (FC) strength as the coupling between M1 and the cingulo-opercular network (CON), which recapitulated the expected inter-effector organization (**Fig. 5b**). We then used canonical correlation analysis (CCA) to derive a multivariate cellular signature associated with this FC profile. The resulting signature, comprising 12 nonredundant cell populations (**Fig. 5d** and **Supplementary Fig. 8b**), closely matched the inter-effector FC map (*r* = 0.872, *P*_smash_ < 0.001; **Fig. 5c**, **d**). We further tested whether this cellular signature generalized beyond M1 by projecting its weights across the cortex and comparing the resulting pattern with inter-effector-to-cortex functional connectivity. The projected cellular pattern closely recapitulated the distributed FC profile (*P*_smash_ < 0.001), with prominent correspondence in the anterior prefrontal cortex, including regions of the CON, and along the borders of visual cortex (**Fig. 5e** and **Supplementary Fig. 8c**). These findings indicate that the cellular signature from M1 inter-effector regions extends beyond the local motor cortex and captures distributed functional organization across the cortex.

### Cellular organization maps onto visual hierarchy and eccentricity

We next examined whether RIDGE-derived cell-type maps captured multiple organizational dimensions within the visual cortex, defined using HCP-MMP1 parcels^35^ (**Fig. 6a** and **Supplementary Fig. 9a**). PCA of the 24 maps identified two dominant spatial components that explained 40.9% and 16.0% of the variance, respectively (**Fig. 6b** and **Supplementary Fig. 9b**). PC1 was weighted most strongly by L4 IT and Chandelier neurons, whereas PC2 was primarily weighted by excitatory neuron types, particularly L2/3 IT and L5 IT neurons (**Fig. 6b**). We then evaluated these components against established features of visual cortical organization. PC1 showed a strong positive association with cortical geodesic distance (rho = 0.73, *P*_smash_ = 0.004; **Fig. 6c**), consistent with the transition from early visual to higher-order regions^36^. PC2 correlated significantly with eccentricity values derived from the HCP-MMP1 atlas (rho = 0.53, *P*_smash_ < 0.001; **Fig. 6d**), consistent with eccentricity-related organization^37^. These results indicate that visual cortical cell-type composition contains separable spatial dimensions associated with both hierarchical processing and eccentricity. At the local scale, we next focused on the ventral temporal cortex (VTC), where cytoarchitectonic subdivisions and category-selective regions provide well-characterized examples of local visual organization^38,39^. To examine their local expression, we evaluated PC1 and PC2 using VTC subdivisions linked to the corresponding organizational dimensions: cytoarchitectonic FG2–FG4 subdivisions reflecting hierarchical differentiation along the ventral visual stream^40^, and category-selective regions reflecting eccentricity-related organization^41^. PC1 increased progressively from FG2 to FG4, paralleling the increase in geodesic distance across cytoarchitectonic subdivisions and extending the hierarchical pattern observed at the system level **(Fig. 6a, e**). In contrast, PC2 showed a U-shaped profile across category-selective regions, consistent with the reported eccentricity-gradient reversal across high-level visual cortex^40–44^. We further projected these regions into a cellular space defined by the dominant contributors to PC1 and PC2, represented by L4 IT and L5 IT neurons. Within this space, FG2, FG3, and FG4 were clearly differentiated, whereas face-, place-, body-, and word-selective regions showed an ordered arrangement consistent with their known functional organization (**Fig. 6f** and **Supplementary Fig. 9c**). Thus, the system-level cellular dimensions were also expressed in local cytoarchitectonic and functional differentiation within the VTC.

**Fig. 6.**
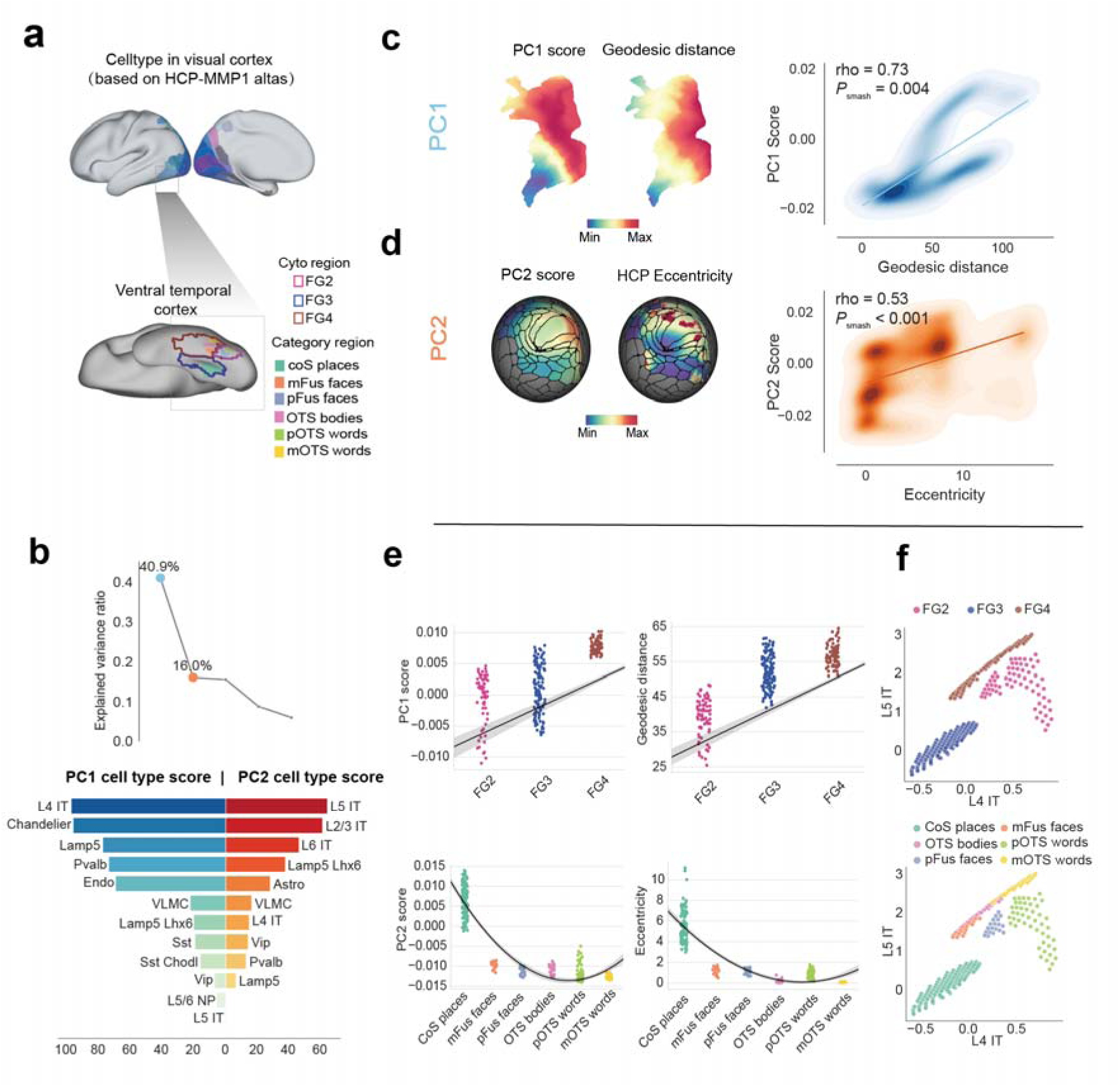
Cell-type maps capture hierarchical and eccentricity-related organization of the visual cortex. **(a)** Visual cortical regions included in the analysis. The visual cortex was defined using 44 areas from the HCP-MMP1 atlas. The magnified view highlights the ventral temporal cortex (VTC), including cytoarchitectonic subdivisions and category-selective regions. See also **Supplementary** Fig. 9a. **(b)** PCA of RIDGE-derived cell-type maps within the visual cortex. PC1 and PC2 explained 40.9% and 16.0% of the variance, respectively. Positive cell-type loadings are shown below. **(c** and **d)** PC1 was positively associated with geodesic distance, whereas PC2 was positively associated with eccentricity. **(e)** Within the VTC, PC1 increased from FG2 to FG4, consistent with geodesic distance (linear fit: PC1, R^2^_adj_ = 0.416; geodesic distance, R^2^ = 0.624), whereas PC2 showed a U-shaped profile across category-selective regions consistent with eccentricity-gradient reversal (quadratic fit: PC2, R^2^_adj_ = 0.847; eccentricity, R^2^_adj_ = 0.795). **(f)** Cytoarchitectonic regions (top) and category-selective regions (bottom) projected into a cellular space defined by the dominant contributors to PC1 and PC2, represented by L4 IT and L5 IT neurons.

## Discussion

RIDGE establishes a reference-guided framework for cortex-wide cellular mapping while preserving regional transcriptional specificity. Across analytical pipelines and independent single-nucleus reference datasets, RIDGE-derived maps showed robust and reproducible cellular organization, recovered established cell-type associations with various cortical features, and recapitulated both broad organizational principles and fine-grained regional specialization across the human cortex. The continuous cellular reference map provides a common substrate for integrating cellular architecture with macroscale brain structure, function and disease.

The regionalized design of RIDGE is motivated by the spatial heterogeneity of cortical cell-type transcriptional profiles^2^. The same cell type can exhibit substantial transcriptional variation across cortical territories, making a spatially invariant reference potentially inadequate for cortex-wide inference^2,15^. Region-matched references therefore provide a means of retaining local transcriptional specialization, whereas holding the cortex-wide transcriptomic anchor fixed provides a common normalization context across regions. This separation allows regional variation to arise primarily from differences in the cellular references rather than from region-specific normalization contexts. Consistent with this rationale, our cross-reference analyses showed greater variability among excitatory populations than among non-neuronal populations, in line with previous evidence for stronger regional transcriptional diversification of cortical excitatory neurons^2,45^. These observations support the view that regionalization in RIDGE reflects biologically meaningful heterogeneity rather than simply increasing model complexity.

The biological relevance of the reconstructed cell-type maps is supported by their convergence with several established dimensions of cortical organization. RIDGE-derived maps recapitulated established relationships involving cortical lamination^17^, evolutionary expansion^18,46–49^, functional hierarchy^1,19,20^, and high-gamma activity^21,50^, indicating that the cell-type distributions retain biologically meaningful variation across distinct organizational domains. Together, these benchmarks point to a broader organizing structure of cortical cellular variation. Recent studies have converged on a robust, evolutionarily conserved principle of cortical specialization, captured spatially by the Pr–Al axis and developmentally by the multinodal induction–exclusion in network development (MIND) model^12,51^. The close correspondence between the dominant cellular pattern reconstructed by RIDGE and the Pr–Al axis, together with its consistency with directly measured single-nucleus cell-type composition, suggests that this pattern reflects a general feature of cortical cellular architecture rather than an artifact of the reconstruction procedure. Comparisons with alternative reference strategies further highlight the importance of balancing regional specificity with cortex-wide coherence. Single-region references could retain aspects of the Pr–Al organization but were more strongly influenced by regionally enriched cellular profiles, whereas cortex-wide robust-marker references may attenuate meaningful regional variation. RIDGE therefore provides a compromise between these two extremes, retaining region-specific transcriptional information while recovering coherent large-scale cortical organization. More broadly, the preservation of the Pr–Al axis indicates that regional transcriptional specialization and cortex-wide continuity represent nested, rather than competing, levels of cellular organization.

This nested organization is particularly relevant in distributed cortical systems, where a shared functional organization coexists with substantial regional cytoarchitectural heterogeneity. The DMN exemplifies this organizational architecture^24^. As an integration-dominant transmodal system, it supports distributed corticocortical integration and exhibits substantial cytoarchitectural heterogeneity across its constituent subregions, reflecting the architectural complexity of this large-scale cortical system^24,25,52^. Within DMN, variation in L2/3 IT, L4 IT, and L5 IT populations was aligned with DMN microstructural differentiation. These intratelencephalic populations contribute substantially to intracortical and corticocortical communication^28,29^, and their regional variation accompanied the transition from highly laminated eulaminate cortex toward less differentiated agranular regions, linking cellular composition to broader cytoarchitectural organization. Such variation may reflect regional differences in the balance between local circuit specialization and distributed corticocortical integration^27,53^. This correspondence nevertheless differed across DMN subregions. The precuneus and middle temporal components showed relatively continuous transitions along the cytoarchitectural axis, whereas the prefrontal component exhibited a spatially interwoven distribution of distinct microstructural types. In the prefrontal DMN, L2/3 IT and L5 IT patterns remained aligned with G1, but neither the histological gradient nor the cellular maps followed the canonical transition from highly laminated to less differentiated cortex. This departure suggests a modular microstructural organization that may support both domain-specific processing and cross-domain integration^28,54–56^. DMN subregions are cellularly differentiated and may support different balances between segregated and integrative processing ^18,57,58^. By resolving both graded shifts in intracortical projection architecture and region-specific departures from the canonical lamination gradient, RIDGE preserves the distinction between shared system-level organization and local specialization within the DMN.

Cellular specialization is also embedded within continuous functional topographies. Within the primary motor cortex (M1), the well-established proximal–distal somatotopic axis provides a clear spatial reference for interpreting the cellular variation resolved by RIDGE. The relative enrichment of L5 ET neurons in proximal territories can be interpreted in light of their role as major corticospinal output neurons providing long-range descending motor signals^33,34^. Oligodendrocyte enrichment in the hand representation accords with the pronounced myelination of this territory, which may support rapid and temporally precise signal transmission during skilled motor control^59,60^. Given their preferential targeting of pyramidal-cell dendrites and their role in shaping local motor-cortical circuits, variation in Sst interneurons may reflect differences in inhibitory regulation across effector representations^61,62^. Considered together, these patterns suggest that the proximal–distal axis is accompanied by coordinated variation across multiple cellular systems rather than by a single dominant cell-type gradient. Beyond classical effector-specific somatotopic organization, inter-effector regions within M1 represent a distinct integrative architecture that links motor territories with distributed extra-motor networks^32^. Their multivariate cellular organization suggests that this integrative architecture may depend on coordinated variation across multiple cell types rather than on any single cellular feature. The correspondence with functionally coupled extra-motor territories suggests a relationship between local cellular specialization and distributed network organization. Consistent with transcriptomic homophily, similarity in regional cellular composition may contribute to preferential functional coupling^50,9,63,64^. Together, these observations provide a cellular perspective on how local motor topography may be embedded within distributed cortical network organization.

Cortical cellular specialization can also vary along multiple separable dimensions. In the visual cortex, these dimensions were associated with hierarchical and eccentricity-based organization^52,65–67^. These cellular dimensions may reflect different balances of feedforward input, local inhibitory regulation, and corticocortical integration. The contributions of L4 IT and Chandelier-type Pvalb interneurons along the hierarchy-related dimensions are compatible with the input-driven architecture of visual cortex. L4 IT neurons constitute a major excitatory population involved in thalamorecipient processing, particularly within the highly differentiated layer 4 that characterizes sensory cortex, whereas Chandelier-type Pvalb interneurons provide rapid axo-axonic inhibition that may support temporally precise sensory processing^17,68–71^. By contrast, the contributions of L2/3 IT and L5 IT populations to the eccentricity-related dimension suggest a greater role for horizontal and long-range corticocortical interactions across visual-field representations^72–74^. Within the VTC, these dimensions were associated with cytoarchitectonic differentiation across FG subdivisions and functional differentiation among category-selective regions ^38,39^. The progression from FG2 to FG4 may reflect a shift in the relative balance between granular, input-associated populations and supragranular or deep-layer populations supporting increasingly integrative processing. The eccentricity-related cellular dimension may likewise contribute to differentiation among category-selective regions, suggesting that foveal and peripheral preferences are accompanied by distinct balances of IT populations^75^. Variation in L2/3 IT organization also offers a possible cellular interpretation of the foveal confluence hypothesis, whereby regions supporting high-acuity object recognition and reading may place greater demands on recurrent intracortical processing, whereas peripherally biased regions may rely more strongly on integration across broader visual contexts^76,77^. Although these interpretations remain tentative, they illustrate how RIDGE-derived cellular dimensions may link system-level visual organization to local cytoarchitectonic and functional specialization.

RIDGE has several limitations. First, the RIDGE-derived maps estimate relative spatial variation in cell-type abundance rather than absolute cellular proportions. These estimates depend on the spatial sampling of the AHBA and the procedures used to generate cortex-wide expression maps, as well as on the anatomical coverage of available single-nucleus references, particularly in regions without directly matched reference samples. Representing cortical cellular diversity using 24 major cell types may also obscure finer regional subtypes and transcriptional states, while cell types represented by relatively few nuclei may be more susceptible to sampling variability and estimation error. Second, RIDGE has so far been evaluated only in the adult cerebral cortex. Its generalizability to other brain structures, developmental stages and disease states remains to be established and will depend on the availability of anatomically and biologically matched reference datasets. Despite these limitations, RIDGE provides a scalable framework for integrating spatially continuous cortical transcriptomic maps with anatomically resolved single-nucleus references while retaining regional cellular specificity. As increasingly comprehensive reference datasets become available, this framework may enable progressively finer and more generalizable characterization of human cortical cellular organization.

## Methods

### RIDGE framework

#### Input data and preprocessing

##### Dense cortical gene-expression maps

The DEMs were used to obtain spatially continuous estimates of cortical gene expression^13^. This resource has been widely adopted in recent studies^78,79^. Derived from the AHBA, probes were reannotated with reference to Arnatkevic iūtė et al. and averaged into gene-level matrices^80^, restricted to the left cerebral cortex due to sparse right-hemisphere sampling (n = 1,304 samples; 20,781 genes). Native sample coordinates were registered to individual surfaces and spatially smoothed using a 20 mm full-width at half-maximum (FWHM) Gaussian kernel. To facilitate cross-donor integration, expression values were z-scored at each vertex and aggregated into the fsLR-32k group space with 29,696 vertices. Processed data are available via the MAGICC project^13^.

For robustness analyses, we additionally used an alternative set of AHBA-derived cortical expression maps generated by Ecker et al.^79^ using a distinct reconstruction pipeline. These maps were resampled to fsLR-32k space for direct comparison with the primary DEMs; details of their generation and preprocessing are provided in the **Supplementary Methods**.

##### Single-nucleus data processing

Two large-scale, multi-region human single-nucleus RNA-sequencing datasets from Siletti et al.^1^ and Jorstad et al.^2^ were accessed through the CELLxGENE Census platform. Analyses were restricted to the cerebral cortex. Low-quality cells (<100 genes) and rare genes (<5 cells) were removed. Following the cellular reference from Siletti et al.^1^, samples with neuronal proportions below 50% were excluded, followed by clusters with poor cortical representation (<10% cortical cells). Next, 3,000 highly variable genes were selected for dimensionality reduction. PCA was performed using 30 components, followed by Harmony correction for dataset and donor effects^81^. Cell identities were subsequently re-annotated into 24 standardized cortical cell types using the label transfer workflow implemented in Seurat^82^. For each target region, label transfer was performed separately against all eight spatial reference regions from Jorstad et al.2. A post-transfer spatial concordance filter was then applied, retaining cells whose assigned identities were supported by the matched regional reference pattern (final n = 1,464,285). Differential-expression analysis was performed using Memento^83^. For each cell type, we compared gene expression in each target cortical region with expression in all other cortical regions. Genes were classified as enriched in the target region if the Memento differential mean expression coefficient exceeded 1 on the natural-log scale and the Benjamini–Hochberg-adjusted *P* value was below 0.05.

#### Multi-region transcriptomic deconvolution

RIDGE integrates region-specific single-nucleus reference profiles with the spatially continuous DEM target to estimate vertex-wise cell-type distributions while maintaining a common transcriptional basis across cortical regions. Single-nucleus profiles and DEMs were first restricted to shared genes. Cell-type distributions were estimated using dtangle^84^, which has previously been applied to deconvolution analyses integrating single-nucleus RNA-sequencing references with microarray-derived tissue expression data^85,86^.

##### Regional reference anchoring

For each Brodmann-defined cortical region, shared genes were retained between the regional single-nucleus reference matrix *S_n_*\* and whole-cortex target matrix M*. To harmonize expression distributions between the regional single-cell reference and the transcriptomic target while establishing a common cortex-wide normalization context, the two matrices were concatenated and jointly normalized. The resulting input matrix was defined as:

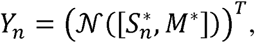

Where N(.) denotes between-sample normalization. The same cortex-wide target M* was used for all N regional references, providing a common normalization context across regions.

##### Marker selection and abundance estimation

Marker genes were identified separately for each region using the jointly normalized input matrix Y_n_. For cell type k in region n, the marker score of gene g was defined as its mean expression in cell type k relative to its mean expression across the remaining cell types:

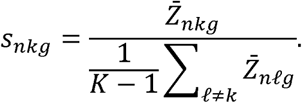

Here, Z_nkg_ denotes the mean expression of gene g in reference cells of type k within Y_n_, and K denotes the total number of cell types. Genes exceeding the cell-type-specific (1 - q) quantile threshold were retained as the regional marker set G_nk_, following the ratio-based marker selection implemented in dtangle. For region-matched deconvolution, these marker sets were applied to the corresponding regional reference and ROI-matched DEM target M_n_. For each cortical vertex V within region n, the pairwise log-ratio quantity between cell types K and t was calculated as:

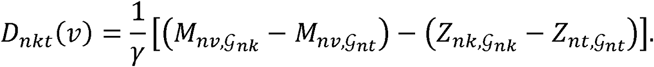

Here, M_nv,Gnk_ denotes the mean target expression at vertex V across marker set G_nk_,Z_nk,Gnk_ denotes the corresponding mean reference expression for cell type k, and is γ the platform adjustment parameter. Cell-type proportions were then estimated using the multivariate logistic transformation:

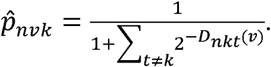

##### Regional reference assignment and cortex-wide integration

For cortical regions with directly matched single-nucleus data, estimates derived from the corresponding regional reference were used for regional estimation. For regions without directly matched single-nucleus samples, surrogate references were selected using a multi-criteria strategy incorporating AHBA transcriptomic similarity, Euclidean proximity and geodesic adjacency. Regional estimates were then integrated across the cortical surface, spatially smoothed using a 6-mm FWHM Gaussian kernel and z-scored to generate continuous vertex-wise distributions of 24 cortical cell types in fsLR-32k space.

### Framework robustness and reference sensitivity

We evaluated RIDGE across four complementary dimensions: (i) robustness to cortical transcriptomic reconstruction, by repeating the complete workflow using alternative AHBA-derived expression maps generated by Ecker et al.^79^; (ii) reproducibility across independent multi-region single-nucleus reference datasets, by comparing RIDGE-derived maps based on the Siletti et al.^1^ and Jorstad et al.^2^ datasets within matched cortical regions; (iii) sensitivity to regional transcriptional context, using systematic cross-regional projection of region-specific references; and (iv) sensitivity to reference strategy, by comparing RIDGE-derived maps with those generated using Lake-derived single-region references and a cross-area robust-marker reference. Detailed procedures are provided in the **Supplementary Methods**, and the transcriptomic datasets and derived resources used in this study are summarized in **Supplementary Table 8**.

### Global-scale evaluation of cell-type maps

#### Prespecified associations with established cortical features

To evaluate the biological plausibility of the RIDGE-derived cell-type maps at the global scale, we aligned the maps to the Schaefer-400 parcellation and tested four prespecified associations between selected cellular measures and established cortical features. These analyses examined: (i) the association between a composite L2/3 IT–L5 IT measure and evolutionary cortical expansion; (ii) the association between L4 IT distribution and layer 4 thickness; (iii) the association between the OPC-to-oligodendrocyte ratio and the principal functional connectivity gradient; and (iv) the association between Pvalb distribution and high-gamma power. The composite L2/3 IT–L5 IT measure was calculated by summing the standardized parcel-wise combined distributions of L2/3 IT and L5 IT neurons. The evolutionary expansion maps, principal functional connectivity gradient, and high-gamma power map were obtained through the neuromaps toolbox^87^. Layer 4 thickness was derived from the BigBrain histological atlas, in which cortical layers were segmented using convolutional neural networks, and was obtained through the BigBrainWarp toolbox^88^.

#### Pr–Al-related organization of cell-type composition

For analyses of the RIDGE-derived cell-type maps, cell-type distributions were summarized according to the HCP-MMP1 parcellation to provide more precise delineation of the cortical territories corresponding to the primary sensory and allocortical reference regions^89^. Based on the parcel-wise cell-type compositions, we calculated the Pr–Al index, where Pr denotes primary sensory regions and Al denotes allocortical/periallocortical regions, following previous studies^12^. V1, S1, and A1 were used as Pr anchors, whereas Pir and LEC/MEC (Ent) were used as Al anchors. For each parcel, Pearson correlation coefficients were calculated between its cell-type composition and those of the Pr anchors as well as the Al anchor parcels. The maximum correlation among Pr anchors and the maximum correlation among Al anchors were retained as Cor_pr_ and Cor_Al_, respectively. Cor_pr_ and Cor_Al_ were then normalized independently by setting the minimum value to 0 and the maximum value to 1. The Pr–Al index was calculated as:

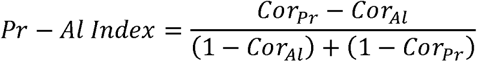

To assess the consistency of the RIDGE-derived Pr–Al organization with directly measured cellular composition, we used processed data from reference^12^. In the human dataset, the correlation between PC1 scores derived from directly measured cell-type composition and the corresponding Pr–Al values was assessed across matched cortical regions, whereas the macaque and marmoset data were included as cross-species references.

To assess the sensitivity of Pr–Al-related organization to reference strategy, we compared the RIDGE multi-region reference strategy with the Lake visual, Lake frontal and cross-area robust-marker reference strategies using spatial topographic measures. Correspondence between the Pr–Al index and the sign-aligned PC1 was quantified across progressively increasing Pr–Al tails using cortical-surface geodesic deviation and Dice overlap. Spatial significance of the geodesic correspondence was assessed using spin permutations, with detailed procedures provided in the **Supplementary Methods**.

### Multivariate associations with neurotransmitter and disease profiles

To further evaluate the cross-modal relevance of the cell-type maps, we examined their multivariate associations with disorder-related cortical abnormalities and neurotransmitter architecture. Cortical abnormality maps comprising standardized effect sizes from more than 21,000 patients across 13 disorders were obtained from the Enhancing NeuroImaging Genetics through Meta-Analysis (ENIGMA) consortium and ENIGMA toolbox^22,90^. Because these maps were defined using the Desikan–Killiany atlas^91^, the cell-type maps were re-aggregated to the same parcellation. For each disorder, we modeled cortical abnormalities as a function of cell-type distributions and used SHapley Additive exPlanations (SHAP) values to quantify the marginal contribution of each cell type^92^. Statistical significance was assessed using spatial permutation testing (n = 1,000). In a separate analysis of neurotransmitter architecture, analyses were restricted to neuronal cell types and receptor-related maps. Receptor density maps covering 19 targets from 26 independent positron emission tomography (PET) studies were obtained from Hansen et al.^23^. We used partial least squares correlation (PLSC) to identify latent variables maximizing shared covariance between neuronal cell-type distributions and receptor architecture, with significance assessed using spatial permutation testing (n = 10,000).

### System-and local-scale analyses of cell-type maps

#### DMN microstructural organization across system and local scales

We examined how the cell-type maps relate to DMN microstructural organization at the system and local scales. At the system scale, the BigBrain primary microstructural gradient (G1) served as a histological reference of cytoarchitectural differentiation. Following Paquola et al.^27^, G1 was derived from the similarity of intracortical staining profiles using diffusion map embedding and obtained through the BigBrainWarp toolbox^88^. Analyses were restricted to the DMN mask defined by the Yeo-7 parcellation, within which vertex-wise correlations were computed between each cell-type map and BigBrain G1. Statistical significance was assessed using spatial permutation testing with FDR correction across cell types. To examine the relationship between cellular organization and network communication, we calculated navigation efficiency using preprocessed group-averaged Human Connectome Project (HCP) structural connectivity matrices parcellated according to the Schaefer-400 atlas and obtained from Paquola et al.^27^. Using greedy routing navigation^30^, mean navigation efficiency (E_nav_) was computed for each DMN parcel in intra-network, inter-network, and global contexts and correlated with the corresponding cell-type maps.

At the local scale, we focused on six DMN subregions, including precuneus, inferior parietal, parahippocampus, middle temporal, inferior frontal, and prefrontal. Local cytoarchitectonic differentiation was characterized using the Von Economo atlas^31^, which classifies cortical areas according to laminar organization from koniocortical to agranular types. For each subregion, vertex-wise BigBrain G1 values and the cell types showing significant system-level associations with G1 were extracted and grouped by Von Economo class (Kon, Eu-III, Eu-II, Eu-I, Dys, and Ag).

#### Motor topography and inter-effector functional organization

We first defined the primary motor cortex (M1) as Area 4 from the HCP-MMP1 atlas^89^. To capture its functional topography, we segmented M1 into foot, inter-effector, hand, and mouth subregions based on the functional localization atlas by Gordon et al.^32^ Based on this segmentation, we conducted two distinct analyses focusing on the proximal-distal axis and inter-effector regions.

For the proximal–distal axis, we quantified cell-type gradients along the canonical ventral–dorsal coordinate. The flattened M1 surface was partitioned into 100 equidistant bins along this coordinate, and representative cell-type values were computed for each bin. The resulting profiles were normalized to the range [-1, 1] and smoothed using a sliding-window median filter (window size = 5) to generate continuous spatial profiles along the motor body map.

For the inter-effector regions, we investigated their SCAN-related functional organization by quantifying functional coupling with the cingulo-opercular network (CON)^32^. Specifically, FC strength between M1 vertices and CON was computed using Fisher’s z-transformed correlations, and mean FC strength was estimated for each vertex to generate a cortical map highlighting inter-effector regions, following the results in Gordon et al.^32^. To identify the multivariate cellular configuration associated with this connectivity pattern, we employed an iterative CCA, with the spatial pattern of FC strength as the target variable (Y) and the spatial distributions of 24 cortical cell types as predictor variables (X). To reduce multicollinearity, the model was refined by recursively pruning the cell type with the lowest absolute weight in each step. The final model was determined by balancing the number of retained features against the correlation coefficient, with significance verified by a spatial permutation test (n = 1,000). Using the negative canonical weights from the selected model, we calculated a cell-type weighted score at each M1 vertex and quantified its spatial correspondence with the inter-effector FC map. The same cell-type weights were subsequently applied to cortical regions outside M1 excluding inter-effector and effector-specific regions, to generate a projected cell-type weighted score map for comparison with inter-effector-to-cortex FC. Spatial correspondence was quantified as the overlap between the top 10% of vertices in the projected cellular signature and observed FC maps, with significance assessed using spatial permutation testing (n = 1,000).

#### Visual hierarchy, eccentricity, and VTC organization

We examined cellular organization in the visual cortex at the system and local scales. At the system scale, the visual cortex was defined by grouping 44 areas from the HCP-MMP1 atlas^89^, following Chen et al.^35^. PCA was applied to the standardized distributions of the 24 cell types across the left-hemisphere visual cortex to summarize their dominant spatial variation. For each component, PCA generated a spatial score map and corresponding cell-type loadings. The resulting components were related to two established dimensions of visual cortical organization: hierarchical processing and retinotopic eccentricity^66,67^. Visual hierarchy was represented by geodesic distance, calculated as the shortest path along the cortical surface from each vertex to the calcarine sulcus. Retinotopic eccentricity was obtained from population receptive field estimates from the HCP 7T dataset^37^ and resampled to fsLR-32k space. Associations between cellular PC scores and the two visual annotations were quantified using Spearman correlation, with significance assessed using spatial permutation testing (n = 1,000).

At the local scale, analyses focused on the ventral temporal cortex (VTC), including cytoarchitectonic areas FG2–FG4 and category-selective regions for places, bodies, faces, and words. These regions were defined using the Rosenke cytoarchitectonic atlas and a functional atlas based on independent fLoc localizer data^40,43^. Linear regression was used to assess variation in geodesic distance and PC1 scores across ordinal cytoarchitectonic ranks, whereas quadratic polynomial regression was used to characterize variation in PC2 across category-selective regions. Multinomial logistic regression was further used to assess whether cytoarchitectonic and category-selective regions could be distinguished using the cell types with the largest absolute loadings on PC1 and PC2, respectively. Model performance was evaluated across 1,000 stratified random splits using 70% of the data for training and 30% for testing, with nested hyperparameter tuning and class balancing. Classification performance was quantified using the macro-averaged F1 score.

### Quantification and statistical analysis

All quantification and statistical analyses were performed in Python and R. Specific details regarding individual association analyses are provided in the corresponding method sections above. To account for spatial autocorrelation, two spatial permutation frameworks were used according to the spatial coverage of the analysis. For whole-cortex parcellated analyses, significance was assessed using spherical spin tests^93^ (denoted as *P*_spin_). For region-specific analyses restricted to spatially discontinuous cortical masks, primarily within the DMN, visual, and motor regions, significance was assessed using null models generated with BrainSMASH^94^ (denoted as *P*_smash_). Statistical significance was defined as *P* < 0.05 with FDR correction for multiple comparisons, as specified above.

## Data availability

The Allen Human Brain Atlas is available at http://human.brain-map.org. Processed dense cortical expression maps are available at https://github.com/kwagstyl/MAGICC and https://gin.g-node.org/sphache/DecodeGABAData. The cortical single-nucleus RNA-sequencing datasets from Siletti et al.^1^ and Jorstad et al.^2^ are publicly available through the CELLxGENE Census at https://cellxgene.cziscience.com. The Lake et al.^95^ visual and frontal cortex single-nucleus RNA-sequencing data are publicly available from the Gene Expression Omnibus under accession **GSE97930**. Processed cell-type composition data for humans, macaques, and marmosets were obtained from Huang et al.^12^ and are available through MCCSTA https://db.cngb.org/stomics/mccsta/. BigBrain is available at https://bigbrainproject.org, and processed BigBrain-derived data are available at https://github.com/caseypaquola/BigBrainWarp. Human Connectome Project datasets, including the 7 Tesla retinotopy dataset, are available at https://db.humanconnectome.org. Neurotransmitter and other multimodal cortical maps are available through NeuroMaps at https://github.com/netneurolab/neuromaps. Cortical abnormality maps from the ENIGMA Consortium are available through the ENIGMA Toolbox at https://github.com/MICA-MNI/ENIGMA. The preprocessed cortical cell-type maps generated in this study are available at https://github.com/WorKLei/RIDGE.

## Code availability

The code used for data analysis in this study is available on GitHub: https://github.com/WorKLei/RIDGE. Publicly available tools and packages used in this study include: (i) brain visualization tools, including Connectome Workbench 2.0 (https://www.humanconnectome.org/software/get-connectome-workbench), Surfplot (https://github.com/danjgale/surfplot?tab=readme-ov-file), and netneurotools (https://github.com/netneurolab/netneurotools); (ii) snRNA-seq data processing and integration, including Scanpy (https://github.com/scverse/scanpy) and Seurat (https://satijalab.org/seurat/); cell-type deconvolution, using dtangle (https://gjhunt.github.io/dtangle/); differential expression analysis, using Memento (https://github.com/mincheoly/memento-cxg); and (iii) statistical testing, including spatial permutation testing using Neuromaps (https://github.com/netneurolab/neuromaps) and BrainSMASH (https://github.com/murraylab/brainsmash), as well as model interpretation using SHAP (https://github.com/shap/shap).

## Acknowledgments

This work was supported by the Natural Science Foundation of China (grants 82425024, 32541013, 82501807 and 82372049) and Brain Science and Brain-like Intelligence Technology - National Science and Technology Major Project (2021ZD0200500).

## Author contributions

B.L. and M.W. led the project. W.L., B.L., and M.W. conceived the study and designed the experiments. W.L. conducted the analysis, with support from X.T., C.D., J.L., X.Z., X.D., L.D., R.Y., Y.H., Y.Y., C.Y., and L.Q. Data preprocessing was completed by W.L. and C.D. M.W. and X.T. provided critical suggestions for the work. J.L., X.D., and A.L. provided feedback on the analysis. W.L., S.L., and B.L. prepared the figures and drafted the manuscript.

## Supplemental information

Supplementary Information is available for this paper and includes Supplementary Methods, Supplementary Figs. 1–9, and Supplementary Tables 1–8. Supplementary Tables 1 and 3–7 are provided as separate Excel files.

