## Supplemental Information for "A spatially continuous cell-type atlas reveals multiscale organization of the human cortex"

**This Supplementary Information includes:**

**Supplementary Methods**

1. Technical robustness and reference sensitivity analyses of RIDGE

2. Tail-matched geodesic deviation and spatial overlap

3. Navigation efficiency analysis

**Supplementary Tables**

1*. Cortical tissue dissections used to construct the multi-region reference

2. Classification and annotation of the 24 cortical cell types

3*. Cell-type harmonization between Siletti and Jorstad datasets

4*. Region-specific differentially expressed genes across cortical regions

5*. Lake cell-type label transfer

6*. Cross-area robust marker genes

7*. Neurotransmitter and cortical abnormality datasets used in cross-modal analyses

8. Summary of transcriptomic resources used in this study

* Provided in separate Excel files.

**Supplementary Figures**

1. Processing of single-nucleus reference data for regional deconvolution

2. Robustness and reproducibility analyses of cell-type distribution estimates

3. Regional reference sensitivity of cell-type maps

4. Global principal axis of cortical cell-type composition and its association with the Pr–Al index

5. Topographic correspondence of Pr–Al-related cellular organization across reference strategies

6. Cell-type topography relates to cortical vulnerability and receptor architecture

7. Cell-type maps align with microstructural and cytoarchitectural axes in DMN

8. Cell-type organization and feature selection in the motor cortex

9. Cell-type principal components and classification in the visual system

**Supplementary Methods**

**Technical robustness and reference sensitivity analyses of RIDGE**

**Alternative cortical transcriptomic mapping.** To assess the robustness of RIDGE to alternative cortical transcriptomic mapping, we additionally used spatially dense AHBA-derived expression maps generated by Ecker et al.^1^ using a distinct processing pipeline. The AHBA data were processed with abagen, restricted to cortical samples, normalized using a robust sigmoid transformation, and mapped to the FreeSurfer fsaverage6 surface. To increase spatial coverage, samples were mirrored across hemispheres, and expression values were spatially interpolated using Gaussian Process Regression (ordinary kriging) based on samples within a 40-mm geodesic neighborhood, thereby accounting for cortical folding. This procedure yielded vertex-wise expression maps for 15,633 genes, which were subsequently smoothed within a 5-mm vertex neighborhood. The resulting maps were resampled from fsaverage6 to fsLR-32k space and processed using the same RIDGE workflow as the primary dense expression maps (DEMs) (**Supplementary Fig. 2a**).

**Cross-dataset reproducibility analysis.** To assess the reproducibility of RIDGE-derived cell-type maps across independent multi-region single-nucleus reference datasets, we separately performed deconvolution using the Siletti et al.^2^ and Jorstad et al.^3^ references for cortical regions represented in both datasets. Cell-type annotations were harmonized to the common set of 24 cortical cell types, and each reference dataset was processed using the same cortex-wide DEM target and deconvolution workflow. For each matched cortical region and cell type, spatial reproducibility was quantified as the Pearson correlation between vertex-wise distributions estimated from the two reference datasets. These correlations were used to evaluate the consistency of spatial cell-type patterns across independently acquired single-nucleus references **(Supplementary Fig. 2b).**

**Cross-regional projection analysis.** To assess the sensitivity of RIDGE-derived cell-type maps to regional transcriptional context, each region-specific single-nucleus reference was independently projected onto the full cortex-wide transcriptomic target matrix, rather than being restricted to its anatomically matched cortical region. This procedure generated a separate cortex-wide cell-type map for each regional reference. For each cell type, spatial correspondence was quantified between maps estimated using different regional references, providing a measure of cross-reference consistency. Regional transcriptional divergence between the corresponding reference regions was quantified using the number of differentially expressed genes identified for the same cell type across regions. We then examined whether greater regional transcriptional divergence was associated with lower spatial consistency of the resulting cell-type maps (**Supplementary Fig. 3a–c**).

**Alternative reference strategies.** To evaluate the sensitivity of cell-type estimation to reference configuration, we implemented two alternative reference strategies. First, single-region references were constructed from the Lake et al.^4^ visual and frontal cortex datasets. The Lake expression matrices were converted to Seurat objects, after which low-quality cells and rare genes were removed using the same filtering criteria applied to the primary single-nucleus datasets, followed by normalization and feature matching. Cell-type annotations in each Lake dataset were harmonized to the common set of 24 cortical cell types using Seurat label transfer with the anatomically corresponding Jorstad reference dataset (V1C for visual cortex and dorsolateral frontal cortex for frontal cortex). Because several cell types were not assigned to cells in the Lake datasets after label transfer, the resulting visual and frontal references differed in cell-type coverage (**Supplementary Table 5**). Each Lake-derived reference was then applied independently to the full cortex-wide transcriptomic target matrix, thereby generating cortex-wide cell-type estimates from a single regional reference without region-wise reference matching or cortex-wide integration. Second, following the strategy used previously^5^, we constructed a cross-area robust-marker reference using published marker genes identified as robust across cortical areas in the Jorstad dataset (**Supplementary Table 6**). This reference was likewise applied independently to the full cortical target matrix. Cell-type maps generated using all alternative reference strategies underwent the same downstream procedures as the RIDGE-derived maps, allowing direct comparison across reference configurations.

**Tail-matched geodesic deviation and spatial overlap (related to Supplementary Fig. 5)**

We assessed whether PC1-derived cellular organization recapitulated the spatial topology of the Pr–Al axis using tail-matched geodesic deviation and spatial overlap. Analyses were performed on the left-hemisphere HCP-MMP1 atlas comprising 180 cortical parcels. Parcel-wise Pr–Al index and PC1 values were assigned to fsLR-32k surface vertices using the HCP-MMP1 label file, such that each selected parcel set was represented by the union of its constituent surface vertices.

For each map, parcels were ranked from Al-like to Pr-like values. At each Pr–Al tail threshold , the upper and lower tails were defined as Pr-like and Al-like regions, respectively. Thresholds of 5%, 10%, 15%, and 20% were examined. Corresponding percentile intervals of the Pr–Al index and sign-aligned PC1 were compared separately, such that spatial correspondence was evaluated only between regions occupying the same rank interval.

For two rank-matched parcel sets $A$and $B$, geodesic deviation was quantified on the fsLR-32k midthickness surface using symmetric vertex-wise nearest-neighbor distances:

$$D\left( A,B \right)=\frac{1}{2}\left[ \frac{1}{\left| V\left( A \right) \right|}\sum_{a\in V\left( A \right)} \min_{b\in V\left( B \right)} d_{g}\left( a,b \right)+\frac{1}{\left| V\left( B \right) \right|}\sum_{b\in V\left( B \right)} \min_{a\in V\left( A \right)} d_{g}\left( b,a \right) \right]$$

where$V\left( A \right)$ and $V\left( B \right)$denote the surface vertices belonging to parcel sets and , respectively, and denotes cortical-surface geodesic distance. For each tail threshold, mean geodesic deviation was calculated by averaging across the included Pr-like and Al-like rank intervals, whereas cumulative geodesic deviation was calculated as the corresponding sum.

Spatial overlap between each pair of rank-matched Pr–Al and PC1 regions was quantified using the Dice coefficient:

$$Dice\left( A,B \right)=\frac{2\left| A\cap B \right|}{\left| A \right|+\left| B \right|}$$

For each tail threshold, mean spatial overlap was calculated by averaging Dice coefficients across the included Pr-like and Al-like rank intervals. Because Dice overlap was calculated separately between corresponding rank intervals rather than between their cumulative unions, this measure captures rank-specific topographic correspondence between PC1-derived and Pr–Al-defined cortical regions. For the spatial permutation analysis, we evaluated upper and lower tail thresholds from 5% to 50% in 5-percentage-point increments. For each reference strategy, we applied 1,000 Vasa spin permutations to its Pr–Al index map and recomputed the mean geodesic deviation from the corresponding PC1 map using the same rank-matching procedure. The observed values were compared with the mean and central 99% interval of the spin-based null distribution at each threshold.

**Navigation efficiency analysis (related to Supplementary Fig. 7)**

Structural navigation was assessed using greedy routing navigation. For each DMN parcel, mean navigation efficiency (E_nav_) was calculated separately for intra-network, inter-network, and global target sets as:

$$E_{\mathrm{nav}}\left( i \right)=\frac{1}{N_{i}}\sum_{j\in T_{i}} \frac{1}{L_{\mathrm{nav}}\left( i,j \right)}$$

where $T_{i}$ denotes the set of target parcels considered for source parcel i, $N_{i}$denotes the number of parcels in$T_{i}$, and $L_{\mathrm{nav}}\left( i,j \right)$ denotes the navigation path length from source parcel $i$ to target parcel $j$. Navigation efficiency was calculated separately for intra-network, inter-network, and global targets and correlated with the corresponding RIDGE-derived cell-type maps.

**Supplementary Tables**

**Supplementary Table 1 | Cortical tissue dissections used to construct the multi-region reference; related to Fig. 1**

The table lists tissue dissection labels, corresponding Brodmann area assignments, sample counts, donor IDs, and source datasets. For tissue dissections not explicitly annotated by Brodmann area in the original datasets, assignments were determined based on the primary anatomical description of each sampled tissue and cross-checked against the supplementary information provided in the original Siletti et al.^2^ and Jorstad et al.^3^ studies. For Brodmann areas represented in both datasets, Siletti-derived reference samples were preferentially used in the RIDGE framework.

**Supplementary Table 2 | Classification and annotation of the 24 cortical cell types; related to Fig. 2.**

| **Cell type** | **Description** | **Class** | **Neighborhood** |
| --- | --- | --- | --- |
| Lamp5 | LAMP5-expressing interneuron | GABAergic | GABAergic |
| Lamp5 Lhx6 | LAMP5- and LHX6-expressing interneuron | GABAergic | GABAergic |
| Pax6 | PAX6-expressing interneuron | GABAergic | GABAergic |
| Sncg | SNCG-like interneuron | GABAergic | GABAergic |
| Vip | VIP-expressing interneuron | GABAergic | GABAergic |
| Chandelier | Chandelier parvalbumin-expressing interneuron | GABAergic | GABAergic |
| Pvalb | Parvalbumin-expressing interneuron | GABAergic | GABAergic |
| Sst | Somatostatin-expressing interneuron | GABAergic | GABAergic |
| Sst Chodl | Somatostatin- and Chodl-expressing interneuron | GABAergic | GABAergic |
| L2/3 IT | Layer 2/3 intratelencephalic-projecting | Glutamatergic | Glutamatergic |
| L4 IT | Layer 4 intratelencephalic-projecting | Glutamatergic | Glutamatergic |
| L5 IT | Layer 5 intratelencephalic-projecting | Glutamatergic | Glutamatergic |
| L6 IT | Layer 6 intratelencephalic-projecting | Glutamatergic | Glutamatergic |
| L6 IT Car3 | Layer 6 intratelencephalic-projecting (Car3-like) | Glutamatergic | Glutamatergic |
| L5 ET | Layer 5 extratelencephalic-projecting | Glutamatergic | Glutamatergic |
| L5/6 NP | Layer 5/6 near-projecting | Glutamatergic | Glutamatergic |
| L6 CT | Layer 6 corticothalamic-projecting | Glutamatergic | Glutamatergic |
| L6b | Layer 6b | Glutamatergic | Glutamatergic |
| Astro | Astrocyte | Non-neuronal | Non-neuronal |
| Endo | Endothelial cell | Non-neuronal | Non-neuronal |
| Micro/PVM | Microglia/Perivascular macrophage | Non-neuronal | Non-neuronal |
| Oligo | Oligodendrocyte | Non-neuronal | Non-neuronal |
| OPC | Oligodendrocyte precursor cell | Non-neuronal | Non-neuronal |
| VLMC | Vascular leptomeningeal cell | Non-neuronal | Non-neuronal |

**Note:** The 24 cortical cell types used in this study were organized into three major classes: glutamatergic excitatory neurons, GABAergic inhibitory interneurons, and non-neuronal cells. Inhibitory cell types were further grouped by major developmental origin (MGE or CGE), whereas excitatory cell types were distinguished primarily by laminar identity and projection targets. Non-neuronal cell types included major glial and vascular-associated populations. This common set of 24 cell types served as the reference classification for RIDGE and downstream analyses. The annotations were adopted from the cortical cell-type framework reported by Jorstad et al.^3^

**Supplementary Table 3 | Cell-type harmonization between Siletti and Jorstad datasets**

**Note:** Region-specific correspondence between Siletti cell classes and the common set of 24 Jorstad cell types. The proportion indicates the relative contribution of each Jorstad cell type to the corresponding Siletti class within each cortical region and sums to 1 for each class–region combination. For BA9 and AnG, Jorstad cell-type proportions are reported directly.

**Supplementary Table 4 | Region-specific differentially expressed genes across cortical regions**

**Note:** Region-enriched genes were identified using Memento^6^. The *de* statistics (*coef*, *se*, *pval*, and *padj*) describe differential mean expression, whereas the corresponding *dv* statistics describe differential expression variability. *roi* and *celltype* indicate the cortical region and cell type, and *pct.1* and *pct.2* indicate the percentages of cells expressing the gene in the target region and comparison group, respectively. Only genes with a differential mean expression coefficient on the natural-log scale (de_coef) greater than 1 and a Benjamini–Hochberg-adjusted *P* value (de_padj) below 0.05 are included.

**Supplementary Table 5 | Lake cell-type label transfer**

**Note:** Cell-type annotations in the Lake frontal and visual cortex datasets were harmonized to the common 24-cell-type classification using Seurat label transfer. The *Summary* sheet reports the numbers of cell types before and after transfer and the total number of transferred cells. The *Frontal* and *Visual* sheets provide the detailed correspondence between original and predicted cell-type labels.

**Supplementary Table 6 | Cross-area robust marker genes**

**Note:** The table lists cross-area robust marker genes derived from the Jorstad dataset and used to construct a cross-area robust-marker reference. *cluster* indicates the 24 cortical cell types, and *robust_genes* lists the corresponding marker genes identified across cortical areas.

**Supplementary Table 7 | Neurotransmitter and cortical abnormality datasets used in cross-modal analyses**

**Note:** The table summarizes the external datasets used for cross-modal analyses. The *PET_Receptors* sheet includes 19 neurotransmitter receptor and transporter targets from 26 PET datasets obtained from Hansen et al.^7^. The *ENIGMA_Disorders* sheet summarizes 13 disorder- and trait-related cortical abnormality maps obtained from the ENIGMA Consortium and ENIGMA Toolbox^8,9^, together with the corresponding sample sizes.

**Supplementary Table 8 | Summary of transcriptomic resources used in this study**

| **Resource** | **Species** | **Source modality** | **Data used in this study** | **Role in this study** |
| --- | --- | --- | --- | --- |
| Allen Human Brain Atlas (AHBA) | Human | Bulk microarray | Cortex-wide gene-expression maps generated using the MAGICC^10^ and Ecker et al.^1^ pipelines | Cortex-wide transcriptomic targets and shared anchors for RIDGE; assessment of robustness to alternative cortical expression-mapping pipelines |
| Siletti et al.^2^ | Human | snRNA-seq | Region-specific cell-type transcriptional reference profiles | Construction of region-specific RIDGE references and cross-dataset reproducibility assessment |
| Jorstad et al.^3^ | Human | snRNA-seq | Region-specific cell-type transcriptional reference profiles | Construction of region-specific RIDGE references and cross-dataset reproducibility assessment |
| Lake et al.^4^ | Human | snDrop-seq | Visual- and frontal-cortex cell-type reference profiles used to construct two alternative single-region references | Reference-strategy sensitivity analysis |
| Huang et al. ^5^ / MCCSTA | Human, macaque, and marmoset | Published cross-species derived measures | Pr–Al-related measures derived from the Siletti et al. human, Chen et al. macaque, and MCCSTA marmoset datasets | Cross-species comparison of Pr–Al-related cortical organization |

**Note:** The MAGICC^10^- and Ecker et al.^1^-derived cortical expression maps were generated from the same underlying AHBA transcriptomic data using alternative cortical expression-mapping pipelines. The Siletti et al.^2^ and Jorstad et al.^3^ datasets were used to construct the primary multi-region cell-type reference, whereas the Lake et al.^4^ visual and frontal datasets were used to construct alternative single-region references. For the cross-species analysis, previously derived Pr–Al-related measures reported by Huang et al.^5^, based on the Siletti et al. human, Chen et al.^11^ macaque, and MCCSTA marmoset datasets, were used rather than reprocessing the underlying transcriptomic data.

**Supplementary Figures**

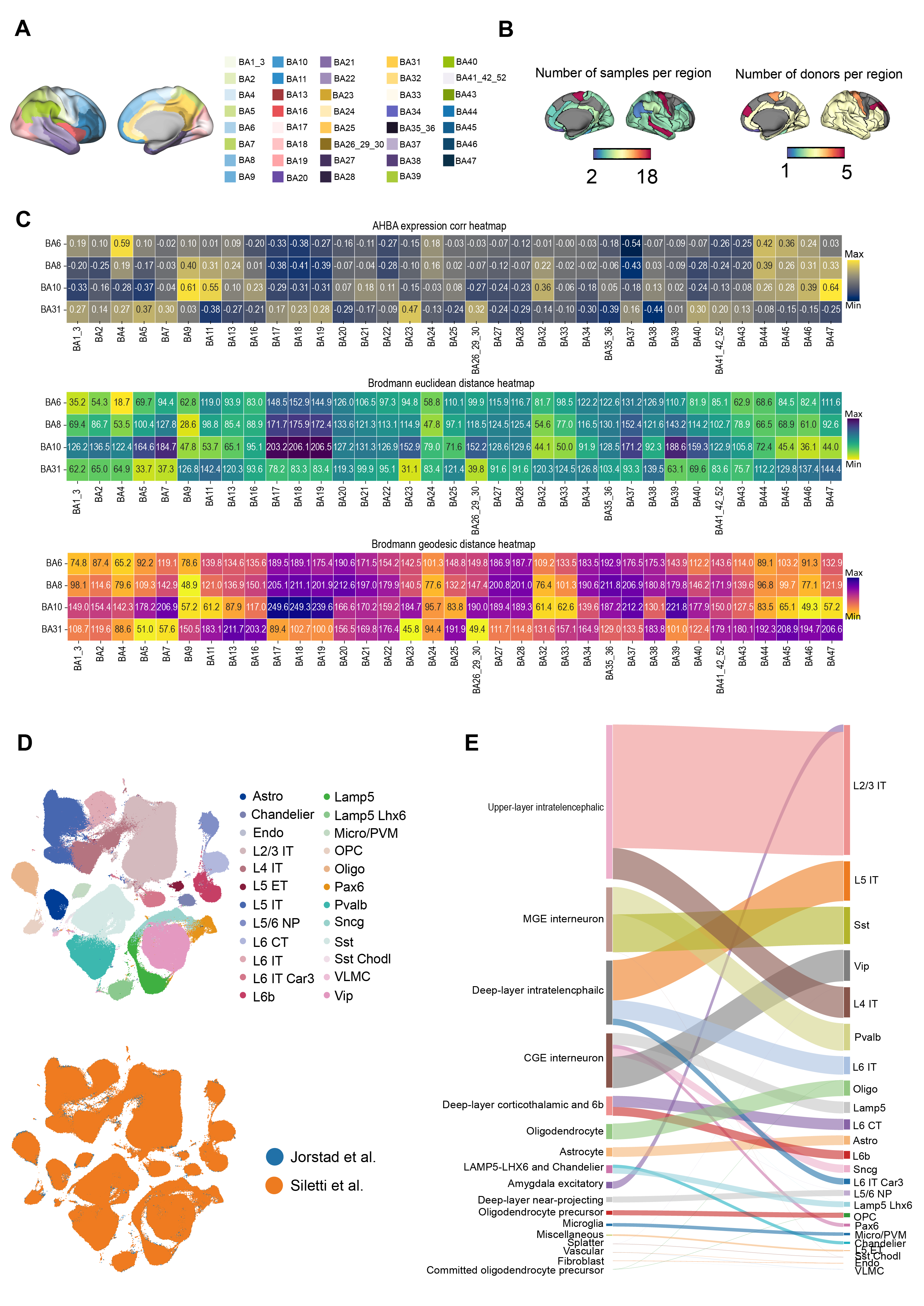

**Supplementary Fig. 1 | Processing of single-nucleus reference data for regional deconvolution,** **related to the “Single-nucleus data processing” section in Methods****.**

**(A)** Brodmann area atlas used for regional deconvolution. Cortical surfaces show the Brodmann area parcellation used to define region-specific transcriptomic references. **(B)** Spatial distribution of available tissue samples and donor coverage across Brodmann areas used for regional transcriptomic deconvolution. **(C)** Multi-criteria substitution strategy for cortical regions lacking direct single-nucleus sampling. Some regions (BA6, BA8, BA10, and BA31) were not represented in the single-nucleus transcriptomic dataset used for regional deconvolution. To ensure comprehensive cortical coverage, substitute datasets were identified based on transcriptomic similarity (AHBA expression correlation), spatial proximity (Euclidean distance), and anatomical adjacency (Brodmann geodesic distance). The heatmaps display pairwise correlations and distances among Brodmann areas, highlighting the regions most similar to each missing area. The cortical surface maps illustrate the distribution of AHBA samples and donors per region, indicating consistency in sampling coverage. Accordingly, BA4 was used as a substitute for BA6, BA9 for BA8, and BA5 together with BA23 for BA10 and BA31, respectively. **(D)** Uniform Manifold Approximation and Projection (UMAP) embedding of the integrated cortical single-nucleus reference dataset. Cells are colored by harmonized cell-type labels (top) and dataset origin (bottom), showing the integrated reference constructed from the Jorstad et al.^3^ and Siletti et al.^2^ datasets. **(E)** Cross-dataset harmonization of cell-type classifications. Sankey plot shows the correspondence between annotation schemes in the Jorstad et al.^3^ and Siletti et al.^2^ datasets.

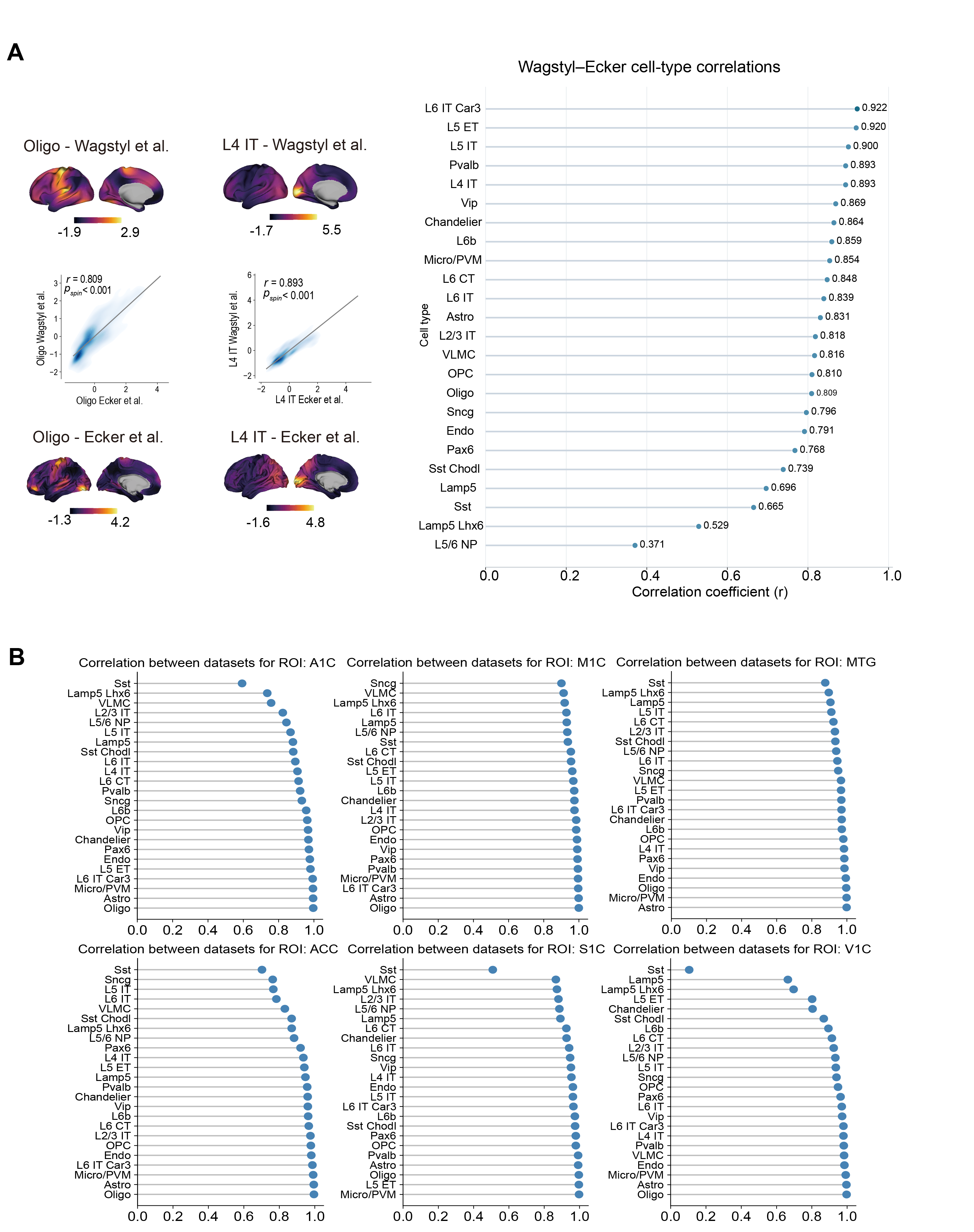

**Supplementary Fig. 2 | Robustness and reproducibility analyses of cell-type distribution estimates, related to the “Framework robustness and reference sensitivity” section in Methods. (A)** Cross-pipeline robustness of surface-based cell-type maps. Representative cortical maps derived using the Wagstyl et al.^10^ and Ecker et al.^1^ preprocessing pipelines are shown for Oligo and L4 IT (spin permutations, n = 1,000), illustrating high spatial correspondence across pipelines. Correlation coefficients between cell-type maps generated from the two preprocessing pipelines are summarized for all 24 cell types, demonstrating consistent spatial patterns across pipelines. **(B)** Cross-dataset consistency of RIDGE-derived cell-type distributions. For each ROI (MIC, A24, A1C, V1C, S1C, MTG), Pearson correlations between cell-type distributions estimated using the Siletti et al.^2^ and Jorstad et al.^3^ single-nucleus reference datasets are shown in descending order.

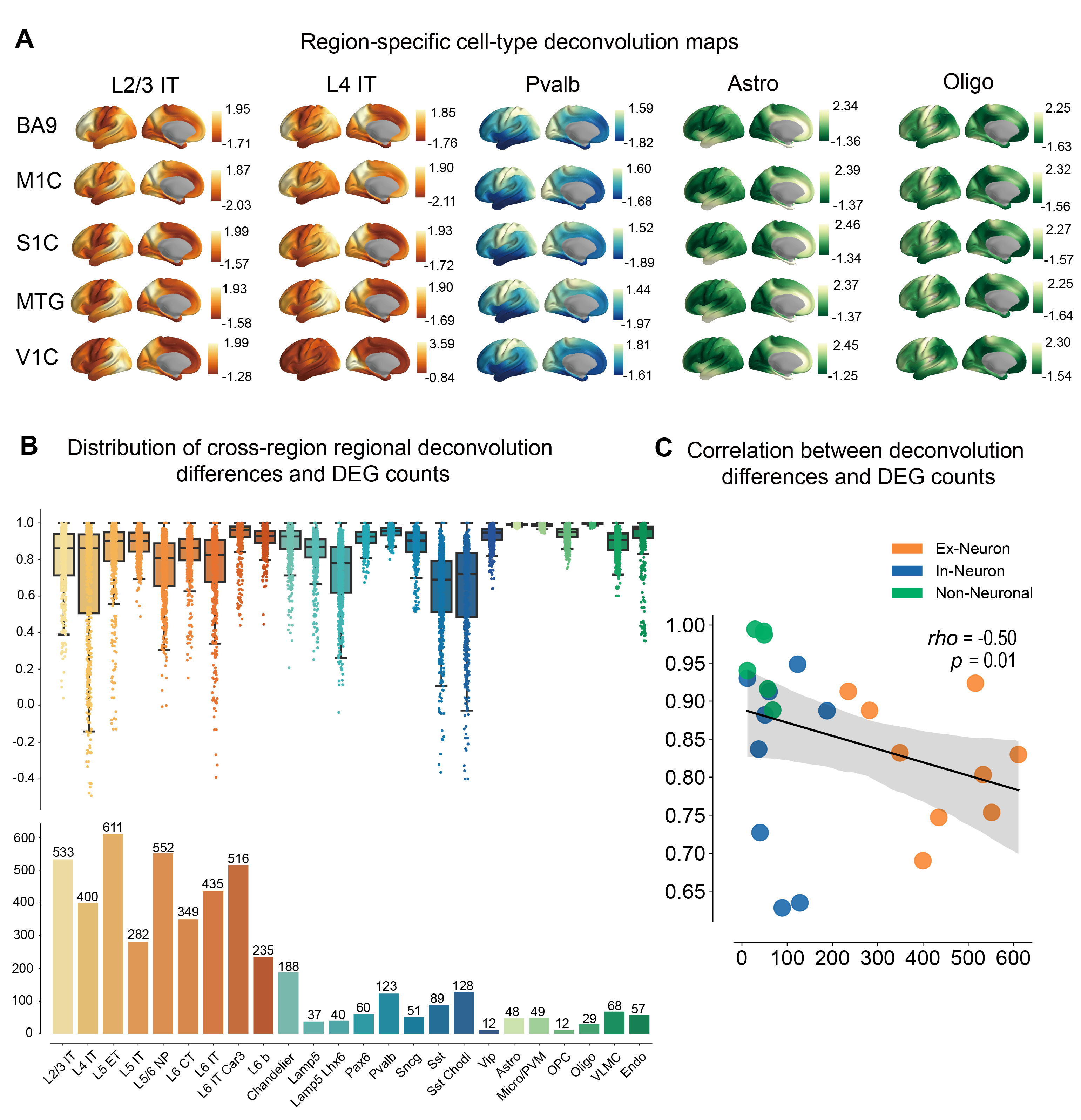

**Supplementary Fig. 3 | Regional reference sensitivity of cell-type maps, related to Fig. 2****. (A)** Cortex-wide cell-type maps estimated using five representative regional references. Each row shows maps estimated using single-nucleus reference signatures from five cortical regions (BA9, M1C, S1C, MTG, and V1C). Excitatory cell types show greater cross-reference variability than non-neuronal cell types, indicating stronger regional specificity. **(B)** Cross-reference variability of cell-type maps and regional transcriptional heterogeneity across cell types. Top, pairwise differences between cell-type maps generated from different regional references. Bottom, the number of region-specific DEGs for each cell type (see Supplementary Table 4). **(C)** Association between cross-reference map similarity and transcriptional heterogeneity across cell types. Each point represents one cell type. Greater regional transcriptional heterogeneity was associated with lower cross-reference map similarity (rho = −0.50, *P* = 0.01). Colors indicate major cell classes.

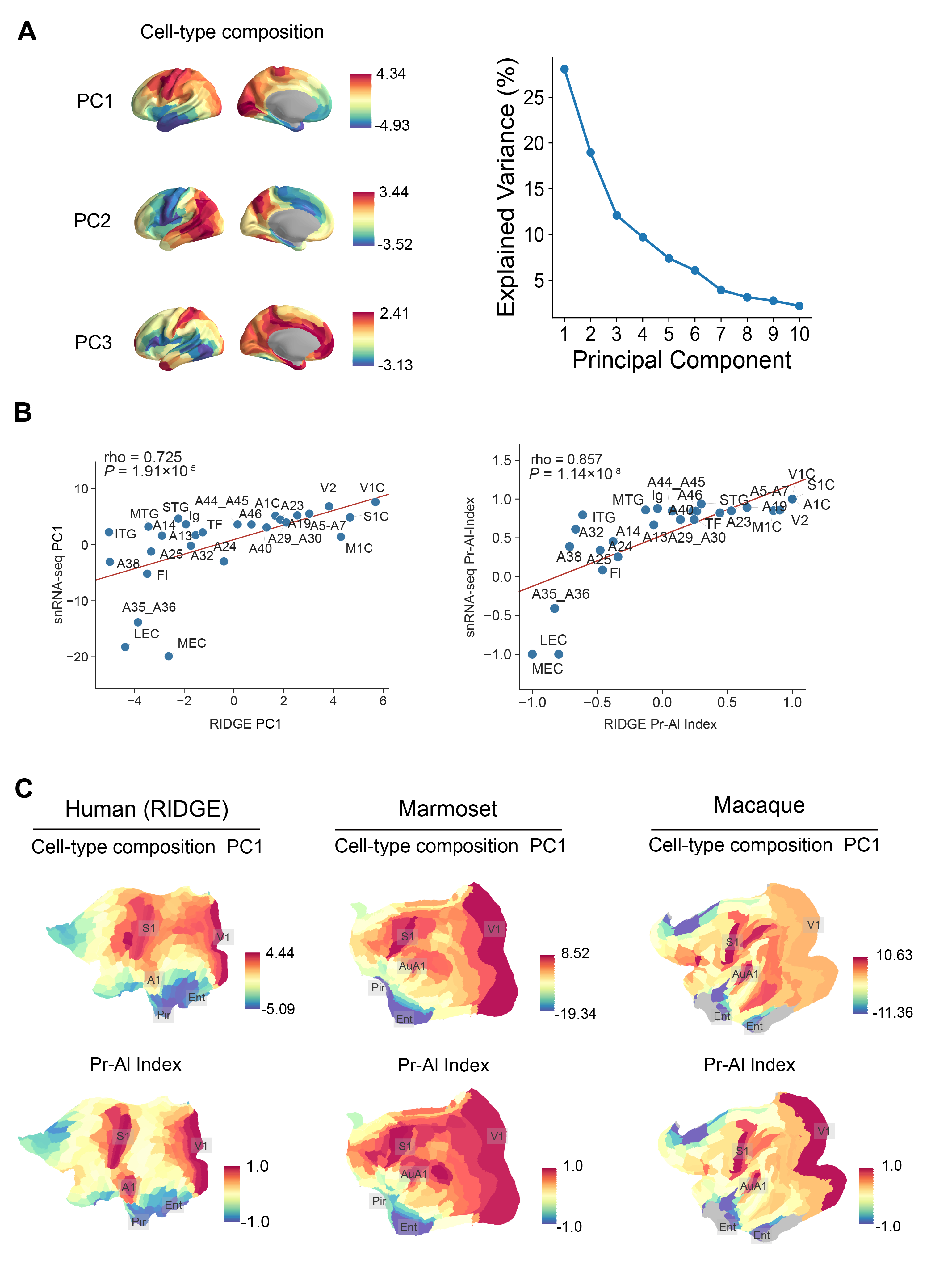

**Supplementary Fig. 4 | Global principal axis of cortical cell-type composition and its association with the Pr–Al index, related to Fig. 3.** **(A)** Principal component analysis of parcel-wise cell-type compositions across the cortex identified a dominant axis of global cellular variation. Left, cortical surface projection of cell-type PC1. Right, scree plot showing the explained variance of the first ten principal components. **(B)** Correspondence with directly measured human single-nucleus cell-type composition. Left, RIDGE-derived PC1 was correlated with PC1 derived from directly measured regional cell-type composition (rho = 0.725, *P* = 1.91 × 10^−5^). Right, spatial distribution of the RIDGE-derived Pr–Al index was correlated with snRNA-seq Pr–Al index (rho = 0.857, *P* = 1.14 × 10^−8^). **(C)** Cross-species comparison of cell-type composition PC1 and the Pr–Al index in humans, marmosets, and macaques.

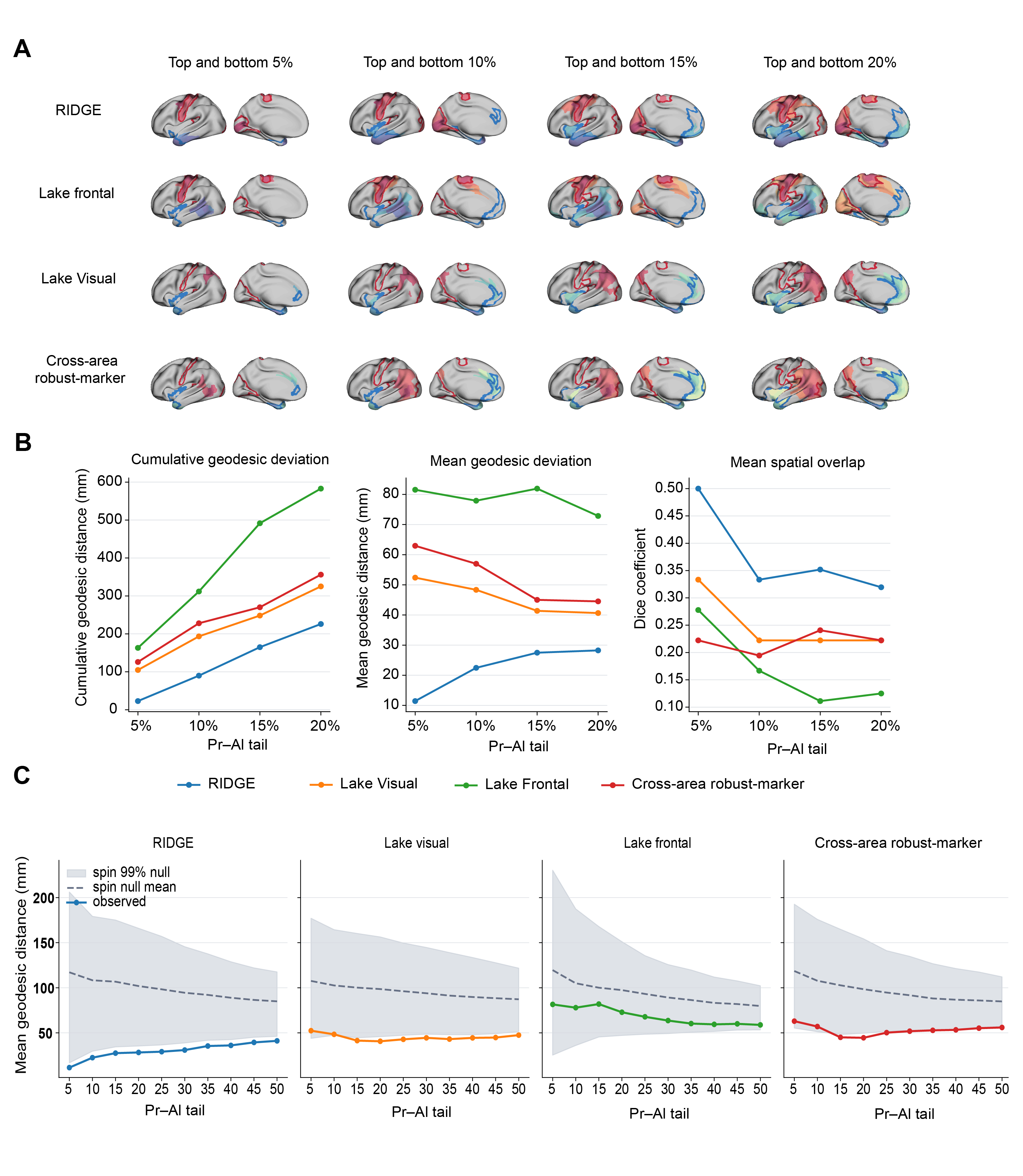

**Supplementary Fig. 5 | Topographic correspondence of Pr–Al-related cellular organization across reference strategies,** **related to Fig. 3. (A)** Spatial correspondence between the Pr–Al index and cell-type composition PC1 at the upper and lower 5%, 10%, 15%, and 20% tails. Four reference strategies were compared: RIDGE, Lake visual, Lake frontal, and cross-area robust-marker. Red and blue contours indicate Pr–Al-defined Pr-like and Al-like regions, respectively, whereas filled regions indicate the corresponding PC1-derived regions for each strategy. **(B)** Quantification of spatial correspondence across increasing Pr–Al tails. The Pr–Al index and PC1 were divided into corresponding percentile intervals from both tails. For each matched percentile interval, cortical-surface geodesic distance and Dice overlap were calculated. Cumulative geodesic deviation represents the sum of geodesic distances across all included percentile intervals, whereas mean geodesic deviation represents their average. Mean spatial overlap represents the average Dice coefficient across matched percentile intervals. Lower geodesic deviation and higher Dice coefficients indicate greater spatial correspondence. **(C)** Spatial permutation assessment of mean geodesic deviation across Pr–Al tails from 5% to 50%. At each tail percentage, spin permutations were used to generate spatial null maps, and mean geodesic deviation was recalculated using the same procedure as for the observed maps. Solid colored lines indicate the observed values for each reference strategy, dashed lines indicate the mean of the spin-based null distribution, and shaded regions indicate the 99% null interval.

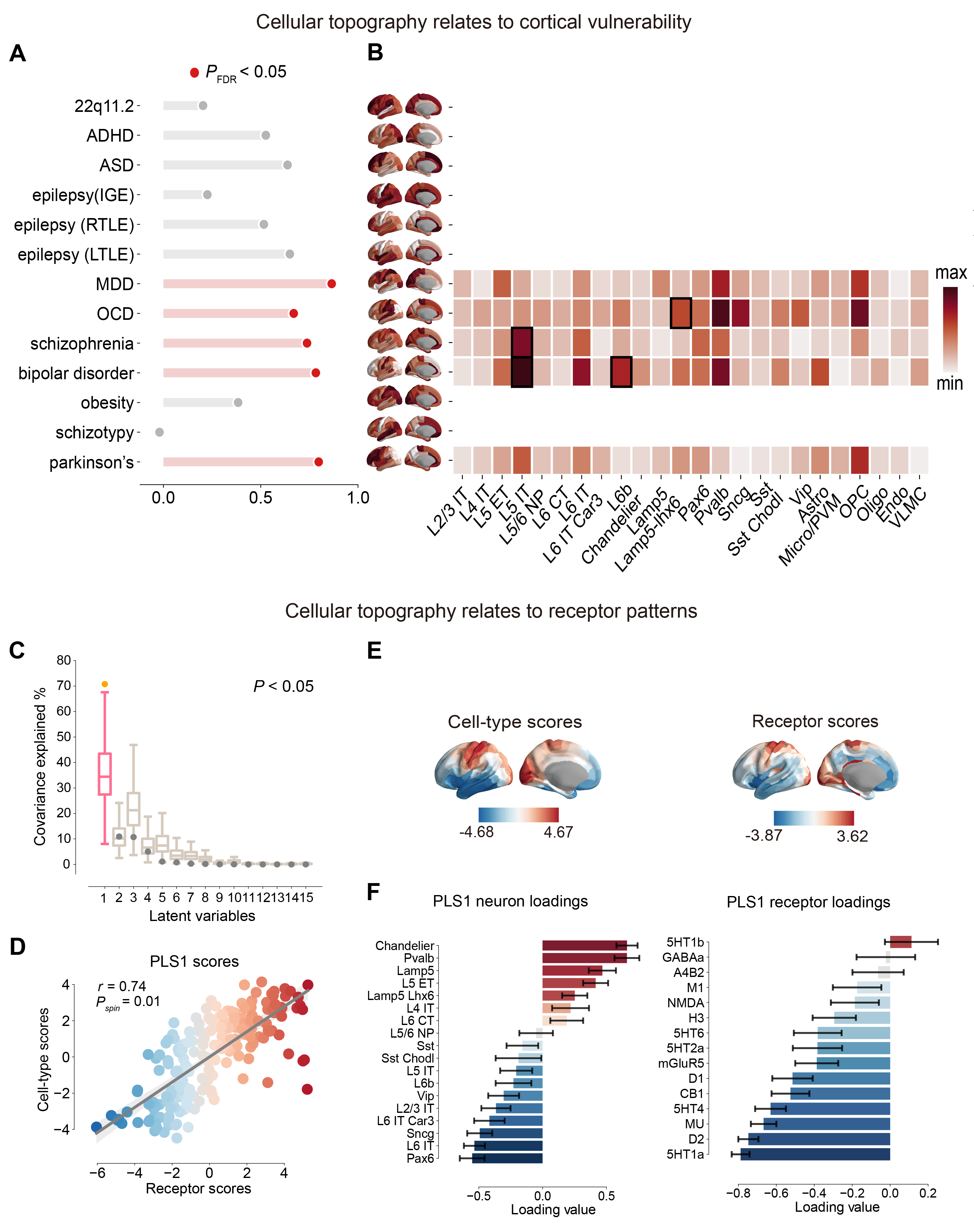

**Supplementary Fig. 6 | Cell-type topography relates to cortical vulnerability and receptor architecture.** **(A)** Multiple linear regression models relating cell-type distributions to cortical abnormalities across 13 disorders from the ENIGMA consortium. Model significance was assessed using spin permutation with FDR correction (n = 1,000, FDR-corrected *P* < 0.05; 22q11.2 deletion R^2^_adj_ = 0.20, *P* = 0.40; ADHD R^2^_adj_ = 0.52, *P* = 0.33; autism R^2^_adj_ = 0.63, *P* = 0.37; epilepsy (IGE) R^2^_adj_ = 0.33, *P* = 0.41; epilepsy (right) R^2^_adj_ = 0.52, *P* = 0.45; epilepsy (left) R^2^_adj_ = 0.65, *P* = 0.33; depression R^2^_adj_ = 0.86, *P* = 0.02; OCD R^2^_adj_ = 0.67, *P* = 0.03; schizophrenia R^2^_adj_ = 0.74, *P* = 0.03; bipolar disorder R^2^_adj_ = 0.78, *P* = 0.01; obesity R^2^_adj_ = 0.38, *P* = 0.34; schizotypy R^2^_adj_ = -0.02, *P* = 0.72; and Parkinson R^2^_adj_ = 0.80, *P* = 0.03). **(B)** Heatmap visualizes the SHAP values for each cell type, reflecting their contribution to the model’s predictions. Higher SHAP values (darker red) indicate greater influence of a given cell type on the model outcome. Black boxes indicate cell-type features with significant spatial contributions (P < 0.05; spin permutations, n = 1,000). **(C)** Partial least squares correlation (PLSC) analysis identified a significant latent variable that explains 71% of the shared spatial variance between neurotransmitter receptor (excitatory and inhibitory receptors) distributions and cell-type maps (excitatory and inhibitory neurons) across the cortex (*P*_spin_ = 0.0004; spin permutations, n = 10,000). The pink box indicates the first latent variable, which reached statistical significance (*P* < 0.05). **(D)** Scatter plot showing the strong correlation between receptor and cell-type scores derived from the first latent variable (*r* = 0.74, *P*_spin_ < 0.01; spin permutations, n = 1,000). **(E)** Receptor and cell-type scores from the first latent variable in Schaefer-400 atlas show a shared topographic gradient linking receptor co-distribution and cell-type maps. **(F)** Receptor and cell-type loadings, defined as Pearson correlations between their cortical distributions and the latent variable scores, indicate each feature’s contribution to the shared spatial pattern; error bars reflect 95% confidence.

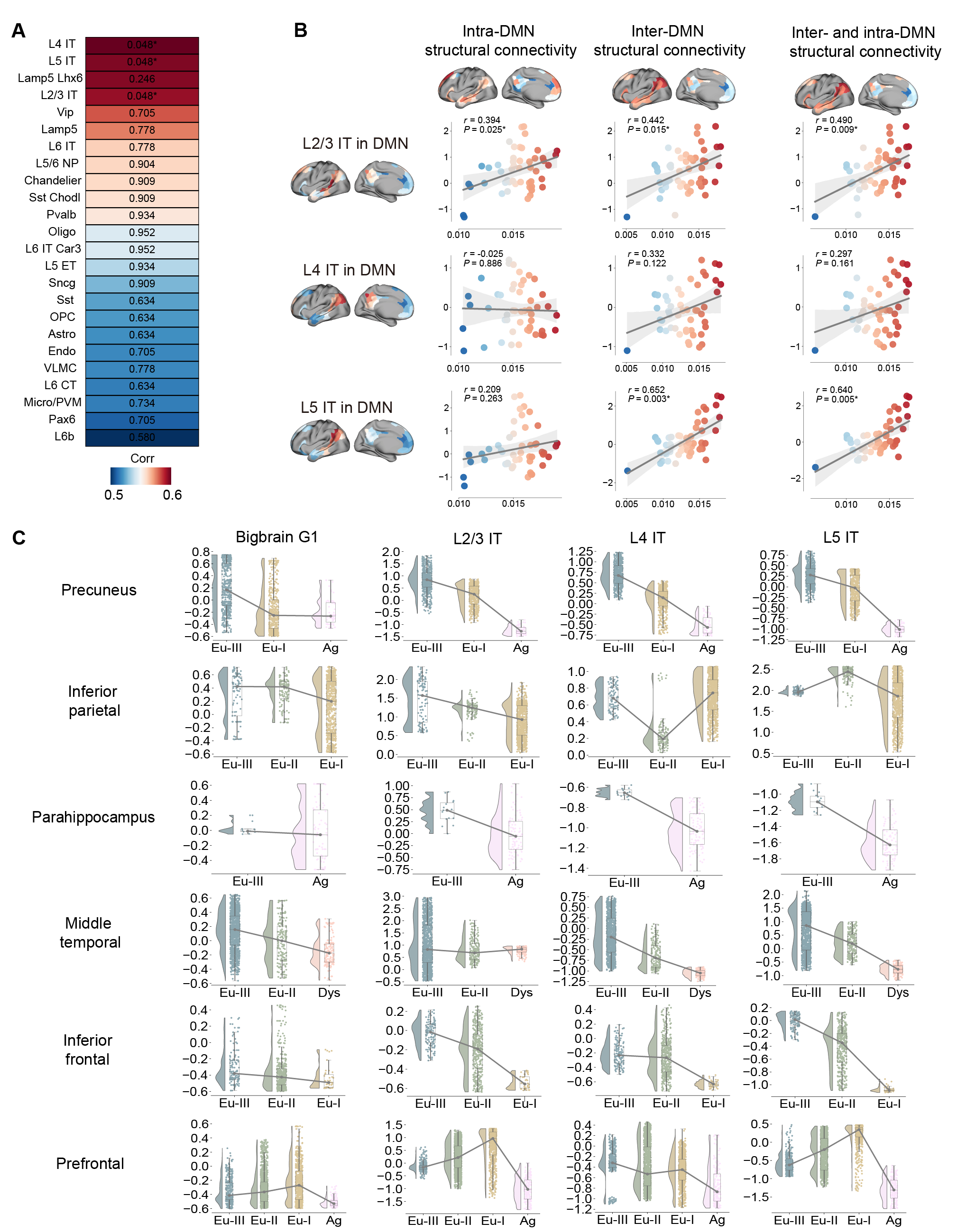

**Supplementary Fig. 7 | Cell-type maps align with microstructural and cytoarchitectural axes in DMN, related to Fig. 4. (A)** Heatmap of *P* values for correlations between cortical cell-type distributions and BigBrain G1 within the DMN, computed using the Schaefer-400 parcellation. Asterisks (“*”) indicate correlations that remain significant after false discovery rate (FDR) correction (FDR-corrected *P*_spin_ *< 0.05;* spin permutation, n = 1,000). **(B)** Spatial distribution of L2/3 IT, L4 IT, and L5 IT within the DMN. Scatter plots show the correlation between cell-type maps and average navigation efficiency (E_nav_) computed for (i) intra-DMN, (ii) inter-DMN, and (iii) combined intra- and inter-DMN (FDR-corrected *P*_spin_ < 0.05; spin permutation, n = 1,000). **(C)** Density profiles show how BigBrain G1 and representative IT cell types (L2/3 IT, L4 IT, and L5 IT) vary across Von Economo cortical types within six DMN subregions (precuneus, inferior parietal, parahippocampal, middle temporal, inferior frontal, and prefrontal cortex).

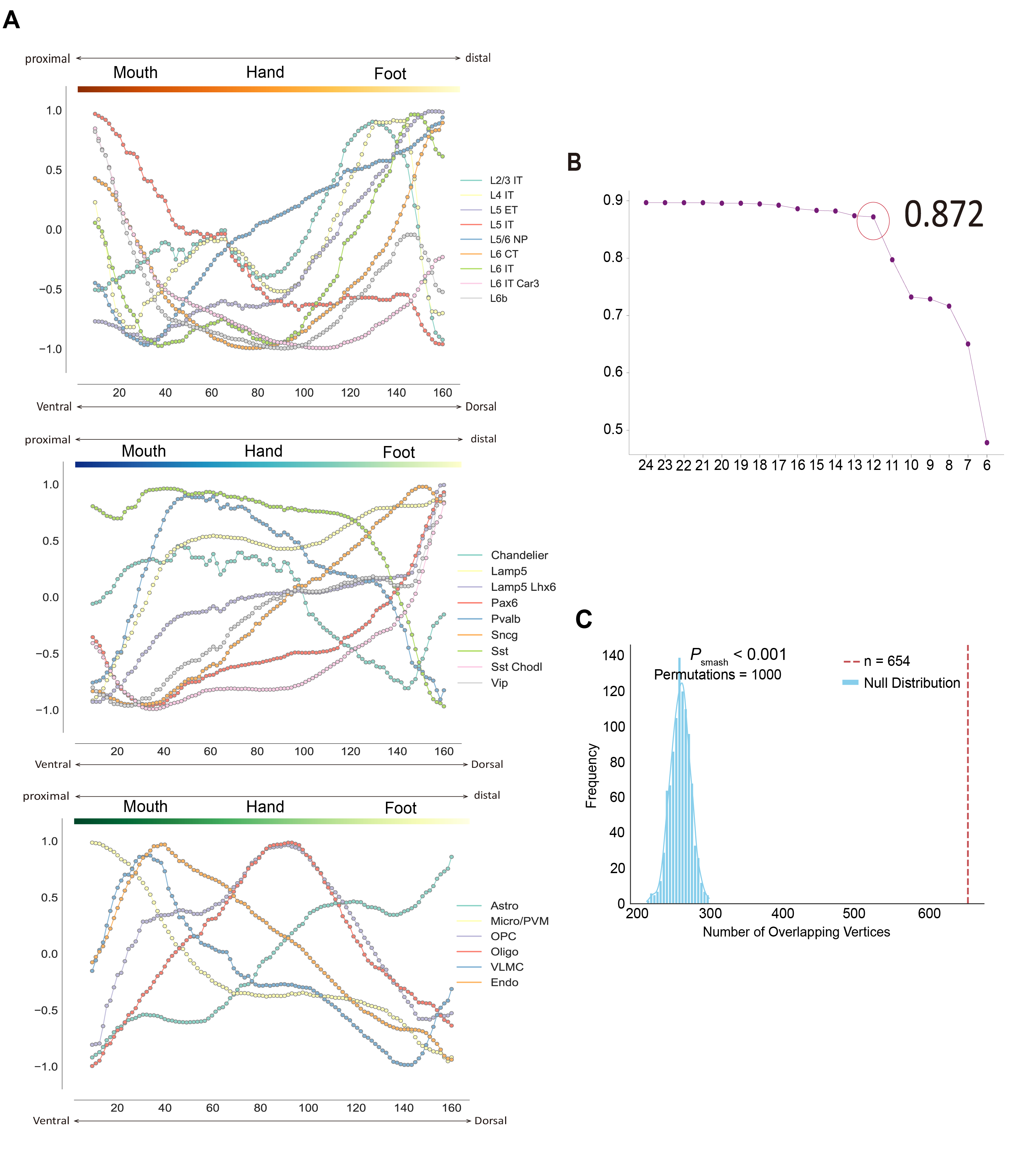

**Supplementary Fig. 8 |** **Cell-type organization and feature selection in the motor cortex, related to Fig. 5. (A)** Cell-type gradients along the proximal–distal axis of primary motor cortex, shown separately for excitatory neurons (top), inhibitory neurons (middle), and non-neuronal cell types (bottom). Each curve represents the normalized distribution of one cell type, scaled to the range [−1, 1] to facilitate comparison across cell classes. The profiles illustrate continuous variation in cell-type composition along the ventral-to-dorsal motor topography. **(B)** Stability analysis of iterative CCA-based feature reduction. The canonical correlation remained stable across successive iterations and dropped sharply at the highlighted step, indicating the optimal stopping point. The retained cell-type set yielded a canonical correlation of 0.872. **(C)** Spatial permutation test for overlap between the cell-type weighted score and inter-effector-to-non-motor functional connectivity strength. The histogram shows the null distribution of top-10% vertex overlap generated from 1,000 spatial permutations. The observed overlap (n = 654) was significantly greater than expected by chance (*P*_smash_ < 0.001).

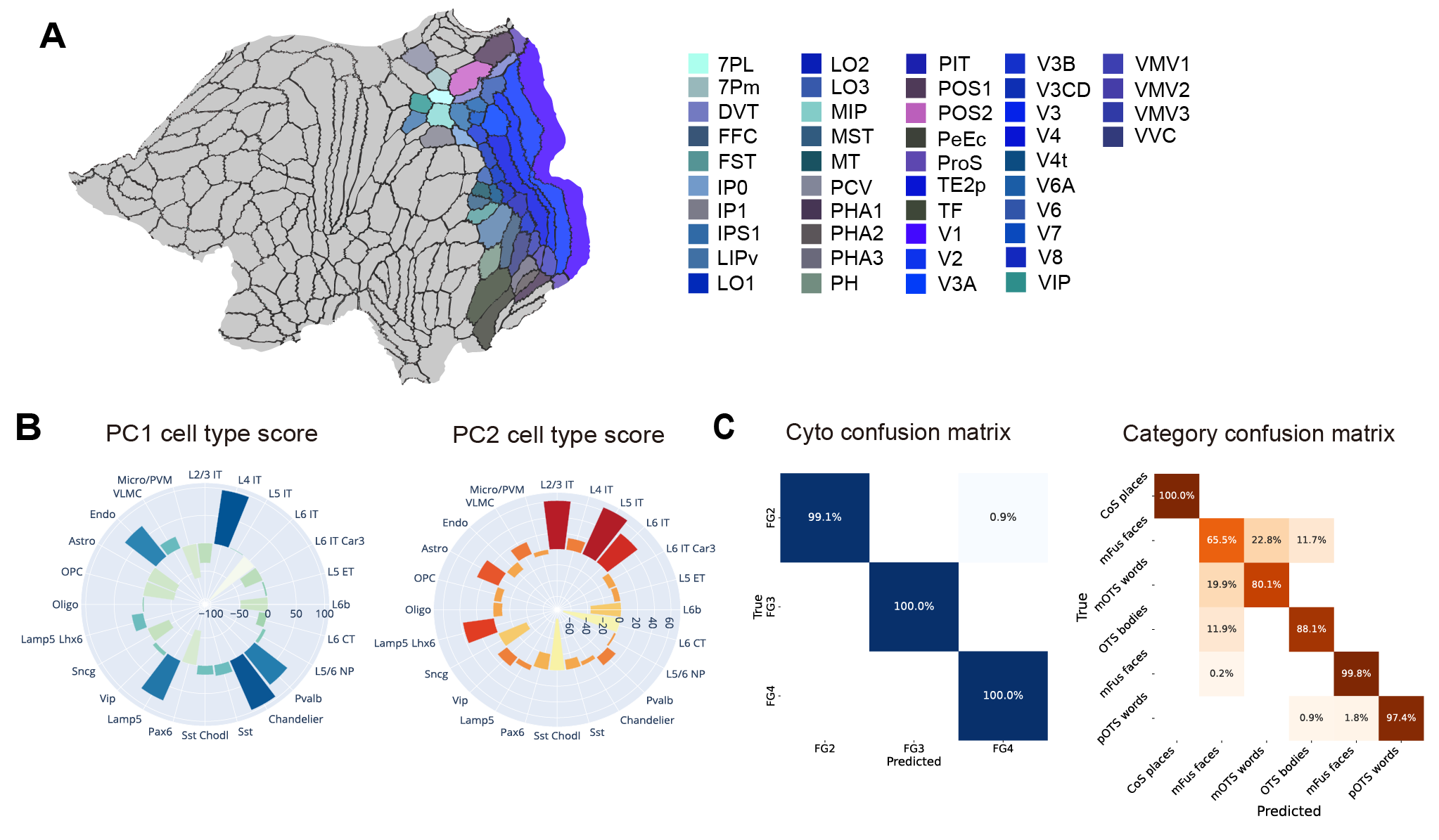

**Supplementary Fig. 9 | Cell-type principal components and classification in the visual system, related to Fig. 6.**

**(A)** Visual cortical regions included in the analysis, defined as 44 areas from the HCP-MMP1 atlas. **(B)** PC1 and PC2 loadings derived from cell-type distributions in the visual cortex, illustrating the principal axes of variation underlying visual cortical cell-type organization. **(C)** Confusion matrices for cytoarchitectonic and category classification using L4 IT and L5 IT, the cell types with the largest absolute loadings on PC1 and PC2, respectively. Left, classification of cytoarchitectonic fields (FG2, FG3, and FG4). Right, classification of category-selective regions (faces, places, bodies, and words). Performance was summarized across 1,000 logistic regression runs.
